# Epithelial-Immune-Stromal Interactions Define Divergent Repair and Fibrosis Pathways After Acute Kidney Injury in Human Renal Transplants

**DOI:** 10.1101/2025.04.30.651080

**Authors:** Shafquat Azim, Thomas Rousselle, Haseeb Zubair, Amol C Shetty, Kellie J Archer, Juliana N Marshall, Ali Rajabi, Caroline M Lara, Sadia Mustofa, Cinthia Drachenberg, Jonathan Bromberg, Madhav Menon, Daniel G Maluf, Enver Akalin, Valeria R Mas

## Abstract

Acute kidney injury (AKI) is a major cause of early graft dysfunction after kidney transplantation, particularly in recipients of high-risk donor kidneys prone to ischemia-reperfusion injury. However, the cellular mechanisms dictating whether injury resolves or progresses to fibrosis remain unclear. This study combines single-nucleus RNA sequencing and imaging mass cytometry (IMC) analysis of human kidney allograft biopsies collected within eight weeks posttransplant, stratified by long-term functional outcomes. Grafts that recovered function were enriched in regenerative proximal tubular (PT) cells co-expressing *PROM1, CD24,* and injury markers, consistent with scattered tubular cells (STCs). In contrast, non-recovering grafts contained a unique subpopulation of transitional proximal tubule cells (tPT4) characterized by dedifferentiation, loss of epithelial identity, and acquisition of fibroblast-like features. Fibroblast trajectory analysis revealed a profibrotic lineage, progressing from stromal progenitors to myofibroblasts, exclusive to nonrecovery grafts. Immune profiling showed divergent macrophage (MΦ) polarization, with reparative MΦ2 cells and regulatory dendritic cell (DC)-like signatures in recovering grafts, versus inflammatory MΦ1 and pro-fibrotic DCs in non-recovery. IMC confirmed spatial colocalization of injured tubules, activated fibroblasts, and immune cells in fibrotic regions, validated in an independent cohort. Functional assays demonstrated that ischemic epithelial injury activated monocyte-derived MΦs with mixed inflammatory/reparative profiles and induced fibroblast-related gene expression, while *PAX8* knockdown impaired epithelial proliferation and promoted pro-inflammatory signaling. These findings reveal epithelial cell plasticity as a central driver of divergent repair outcomes following renal transplant AKI and highlight epithelial–immune–stromal crosstalk as a therapeutic target to promote recovery and prevent chronic graft injury.

**One Sentence Summary:** Single-cell and spatial mapping of human kidney transplants reveal regenerative and fibrotic cell programs across tubular, immune, and stromal compartments that determine whether acute injury resolves or progresses to chronic allograft injury.

## Introduction

Kidney transplantation (KT) remains the most effective treatment for end-stage renal disease (ESRD). However, rising transplantation rates have not kept pace with demand, leading to increased use of high Kidney Donor Profile Index (KDPI) (*1*) grafts, including those from older donors and donation after circulatory death (DCD) donors (*2–6*). These grafts are more susceptible to ischemia-reperfusion injury (IRI), the principal cause of acute kidney injury (AKI) in kidney transplant recipients (KTRs) (*7, 8*). IRI, an inevitable consequence of kidney transplantation, triggers a cascade of stress responses, inflammation, and heightened graft immunogenicity, contributing to delayed graft function (DGF) in 20–50% of deceased donor KTRs (*7–12*).

While AKI is increasingly recognized as a precursor to chronic kidney disease (CKD) in native kidneys (*13–16*), the molecular mechanisms that dictate successful versus maladaptive repair with persistent graft dysfunction following post-transplant AKI remain poorly understood. Human studies of AKI in native kidneys are limited by the invasiveness of biopsy procedures and the absence of longitudinal sampling. In contrast, kidney allografts are routinely biopsied as part of surveillance protocols or to investigate graft dysfunction, offering a unique opportunity to study AKI in a clinically relevant setting. However, recovery from AKI in the transplant context is confounded by donor organ quality, alloimmune activation, and drug-induced nephrotoxicity (*17, 18*), each shaping the injury response and recovery trajectory.

Previous transcriptomic studies have revealed AKI-associated injury and inflammation signatures in human kidney grafts (*19*). Notably, Halloran and colleagues (*20*) identified two AKI subtypes with distinct molecular features and clinical outcomes. However, these bulk-level approaches lack the cellular resolution needed to map how specific epithelial, stromal, and immune compartments coordinate repair or drive fibrosis, processes that are critical to developing any potential therapeutics.

To address this gap, we performed the first comprehensive single-nucleus RNA sequencing (snRNA-seq), and imaging mass cytometry (IMC) analysis of human kidney allograft biopsies collected within eight weeks post-transplant and stratified by functional outcomes and histological evidence of fibrosis progression. Our analysis reveals distinct regenerative and maladaptive cell states, highlighting epithelial plasticity, immune polarization, and fibroblast activation as key determinants of graft recovery or fibrosis. These findings establish the epithelium as a central orchestrator of injury responses, driving either repair or fibrotic remodeling, and uncover novel cellular targets for therapeutic intervention.

## RESULTS

### Patient and sample characteristics

This study evaluated 11 kidney biopsies from unique KTRs using snRNA-seq and IMC, comprising four protocol biopsies from patients with normal allograft function and histology (nKTx) and seven clinically indicated biopsies obtained within eight weeks post-transplant due to acute kidney injury (postKT-AKI). Notably, the nKTx biopsies were collected more than 15 months after transplantation from recipients with stable graft function and normal histological findings, to minimize the confounding effects of early or subclinical injury, thereby representing a transcriptional profile consistent with epithelial homeostasis and resolved injury. The postKT-AKI samples exhibited acute tubular necrosis (ATN) without histological evidence of other acute or chronic allograft injury, as all Banff (*21, 22*) acute and chronic scores were zero. Normal native kidney (NNK, n=3) biopsies from living donors, which experience minimal ischemic injury, were used as controls (**Fig. 1A**).

**Fig 1.**
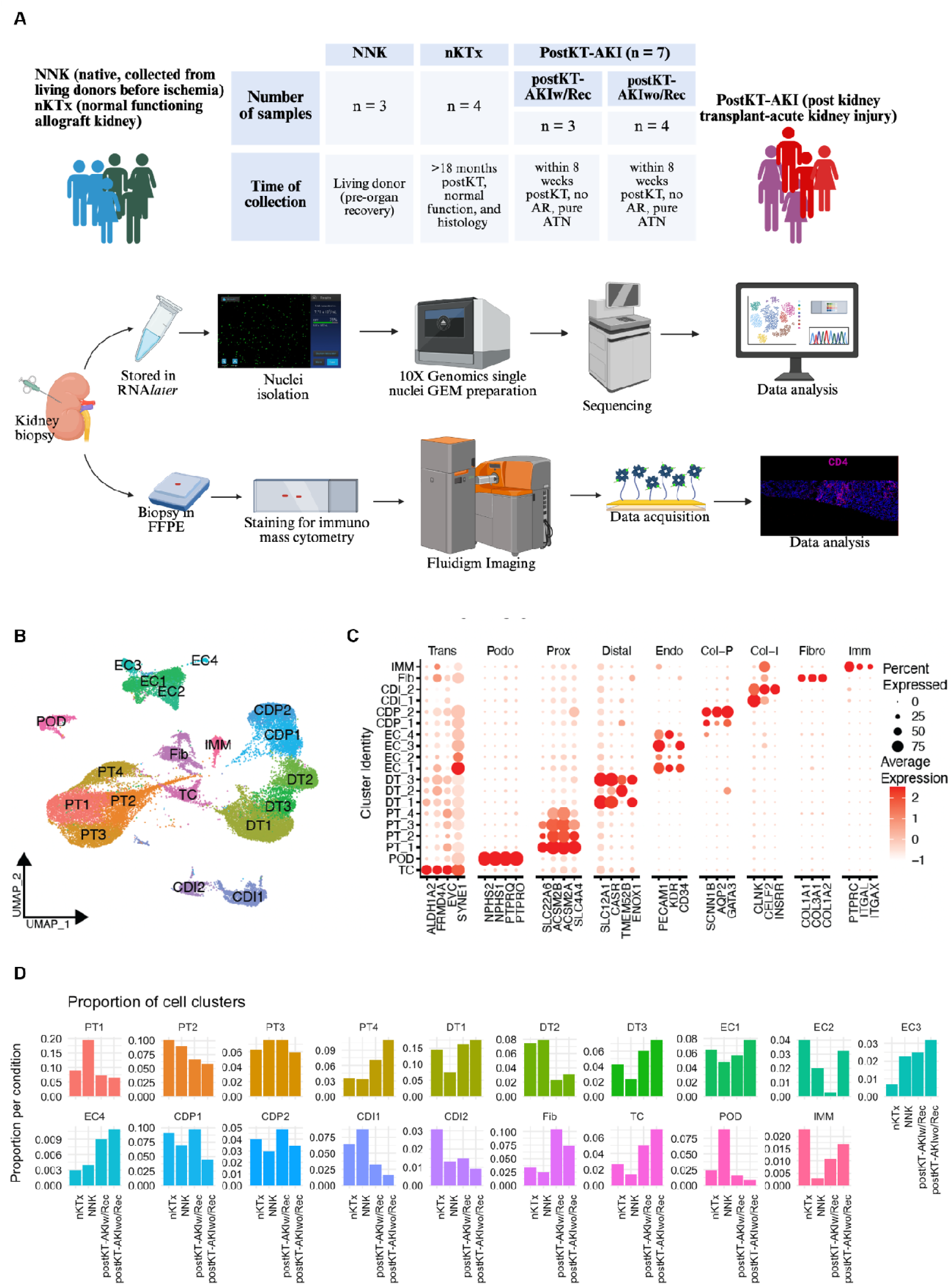
Overview of the study and initial cell cluster analyses. (**A**) Study flowchart demonstrating the study population and procedures used. Biopsies from native (living donor, NNK), normal allograft (nKTx), and graft kidneys with post-kidney transplant acute kidney injury (postKT-AKI) were evaluated using snRNA-seq. Spatial and protein validation was done using IMC. Gene expression changes were studied in immortalized cells using qPCR or bulk-RNA-sequencing. (**B**) UMAP represents cell clusters identified the post-integration of all samples using Seurat. (**C**) Dot plot demonstrates the expression of top gene markers used for the identification of main cell clusters. (**D**) Cell cluster proportions among study kidney groups.

Detailed patient and sample characteristics are presented in **Table 1**. All KTRs received kidneys from deceased donors, including five from DCD and two from donors after brain death (DBD). All patients, except one, received anti-thymocyte globulin induction therapy. One patient with donor-specific anti-HLA antibodies was treated with intravenous immunoglobulin (IVIG) and anti-thymocyte globulin. All KTRs were maintained on a triple immunosuppression regimen consisting of tacrolimus, mycophenolate mofetil, and prednisone. The median cold ischemia time (CIT) in postKT-AKI cases was 31.38 hours (range: 24.56–43.16 hours). Six of the seven developed DGF requiring hemodialysis post-KT. Estimated glomerular filtration rate (eGFR) at 1 week, and at 1-, 6-, 12-, and 24-months post-KT is summarized in **Table 1**. Histological evaluation of NNK samples is shown in **Table 2**.

**Table 1.**
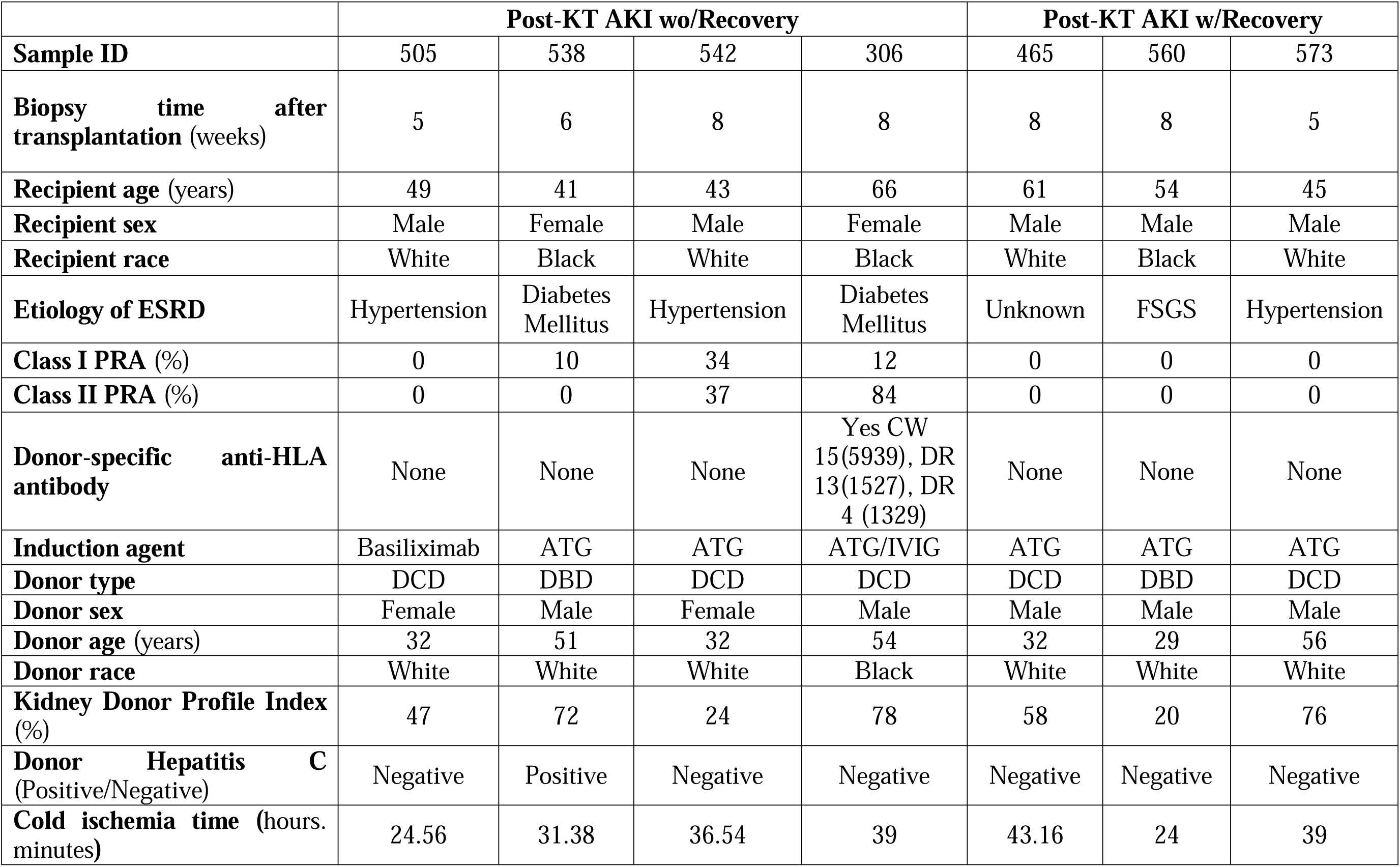
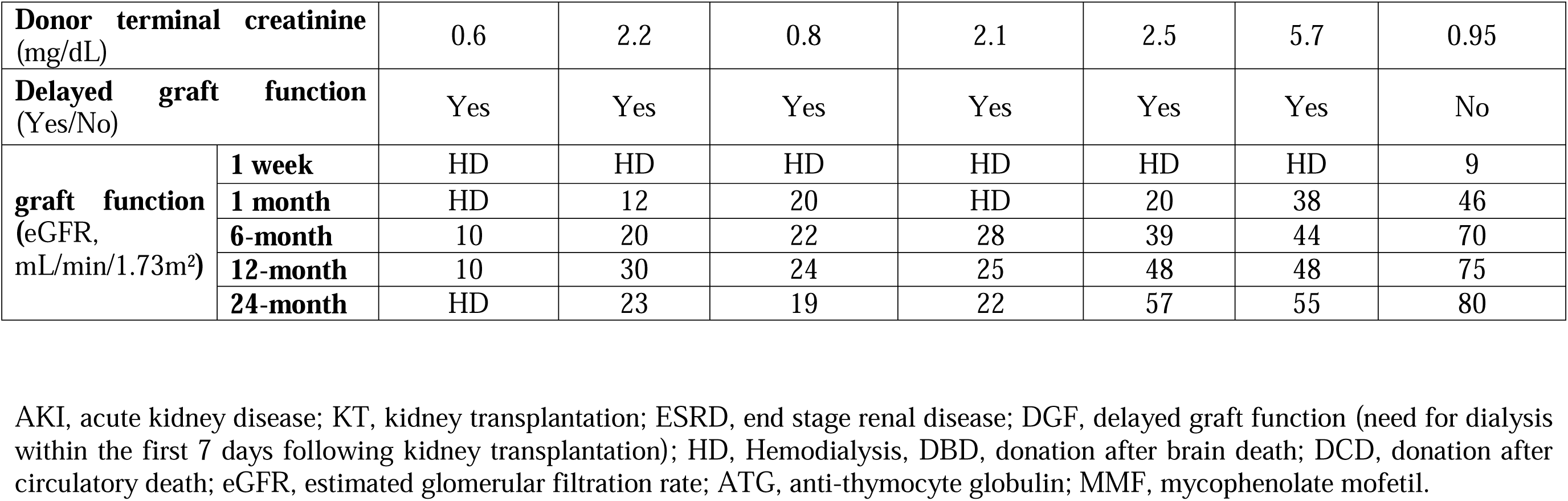
Characteristics of patients and donors from samples included in the study.

|  | **Post-KT AKI wo/Recovery** |  |  |  | **Post-KT AKI w/Recovery** |  |  |
| --- | --- | --- | --- | --- | --- | --- | --- |
| **Sample ID** | 505 | 538 | 542 | 306 | 465 | 560 | 573 |
| **Biopsy time after transplantation (weeks)** | 5 | 6 | 8 | 8 | 8 | 8 | 5 |
| **Recipient age (years)** | 49 | 41 | 43 | 66 | 61 | 54 | 45 |
| **Recipient sex** | Male | Female | Male | Female | Male | Male | Male |
| **Recipient race** | White | Black | White | Black | White | Black | White |
| **Etiology of ESRD** | Hypertension | Diabetes Mellitus | Hypertension | Diabetes Mellitus | Unknown | FSGS | Hypertension |
| **Class I PRA (%)** | 0 | 10 | 34 | 12 | 0 | 0 | 0 |
| **Class II PRA (%)** | 0 | 0 | 37 | 84 | 0 | 0 | 0 |
| **Donor-specific anti-HLA antibody** | None | None | None | Yes CW 15(5939), DR 13(1527), DR 4 (1329) | None | None | None |
| **Induction agent** | Basiliximab | ATG | ATG | ATG/IVIG | ATG | ATG | ATG |
| **Donor type** | DCD | DBD | DCD | DCD | DCD | DBD | DCD |
| **Donor sex** | Female | Male | Female | Male | Male | Male | Male |
| **Donor age (years)** | 32 | 51 | 32 | 54 | 32 | 29 | 56 |
| **Donor race** | White | White | White | Black | White | White | White |
| **Kidney Donor Profile Index (%)** | 47 | 72 | 24 | 78 | 58 | 20 | 76 |
| **Donor Hepatitis C (Positive/Negative)** | Negative | Positive | Negative | Negative | Negative | Negative | Negative |
| **Cold ischemia time (hours. minutes)** | 24.56 | 31.38 | 36.54 | 39 | 43.16 | 24 | 39 |

|  |  |  |  |  |  |  |  |  |
| --- | --- | --- | --- | --- | --- | --- | --- | --- |
| **Donor terminal creatinine**<br>(mg/dL) |  | 0.6 | 2.2 | 0.8 | 2.1 | 2.5 | 5.7 | 0.95 |
| **Delayed graft function**<br>(Yes/No) |  | Yes | Yes | Yes | Yes | Yes | Yes | No |
| **graft function**<br>(eGFR,<br>mL/min/1.73m<sup>2</sup>) | **1 week** | HD | HD | HD | HD | HD | HD | 9 |
|  | **1 month** | HD | 12 | 20 | HD | 20 | 38 | 46 |
|  | **6-month** | 10 | 20 | 22 | 28 | 39 | 44 | 70 |
|  | **12-month** | 10 | 30 | 24 | 25 | 48 | 48 | 75 |
|  | **24-month** | HD | 23 | 19 | 22 | 57 | 55 | 80 |

AKI, acute kidney disease; KT, kidney transplantation; ESRD, end stage renal disease; DGF, delayed graft function (need for dialysis within the first 7 days following kidney transplantation); HD, Hemodialysis, DBD, donation after brain death; DCD, donation after circulatory death; eGFR, estimated glomerular filtration rate; ATG, anti-thymocyte globulin; MMF, mycophenolate mofetil.

**Table 2.** Histological features of nKTx and postKT-AKI samples.

| Banff lesion scores |  | nKTx |  |  |  |
| --- | --- | --- | --- | --- | --- |
|  |  | KUT023 | KUT060 | KUT081 | KUT043 |
|  | Glomerulitis (g) | 0 | 0 | 0 | 0 |
|  | Interstitial inflammation (i) | 0 | 0 | 0 | 0 |
|  | Peritubular capillaritis (ptc) | 0 | 0 | 0 | 0 |
|  | Tubulitis (t) | 0 | 0 | 0 | 0 |
|  | Intimal arteritis (v) | 0 | 0 | 0 | 0 |
| BK virus (SV40 large T antigen) |  | - | - | - | - |
| C4d |  | - | - | - | - |
| Global glomerulosclerosis (%) |  | <5 | 0 | <5 | <5 |

All biopsies were assessed using light (LM), immunofluorescence (IF), and electron microscopy (EM) and reported independently by two transplant pathologists using Banff 2019, update of the Banff 97 classification (21, 22). All biopsies were negative for peritubular capillary staining for complement factor 4 fragment d (C4d) tested by immunofluorescence. All biopsies were negative for BK virus (SV40 large T antigen, immunoperoxidase staining). Abbreviation: nKTx, normal kidney allograft

An independent cohort of paired pre-implantation and 12-week post-transplant biopsies from deceased donor (DD) KTRs was analyzed using microarrays. Patients were stratified as with DGF and non-DGF (postKT-DGF, n=16; postKT-non-DGF, n=29) immediately posttransplant and based on functional outcomes to validate our snRNA-seq analysis, using definitions as described in **Methods and Suppl Inform**. Briefly, KTRs received same triple immunosuppression, included similar distribution of marginal donors, donor age, sex, and age between the two groups. Patient characteristics are described in **Table 3**.

**Table 3:** Characteristics of donors and recipients for samples included in the microarray.

| Characteristics |  | PostKT-nonDGF | PostKT-DGF | *p* value |
| --- | --- | --- | --- | --- |
| **Patients** | N = 45 | n = 29 | n =16 |  |
| **Outcome** | w/Rec | 20 (69.0%) | 10 (62.5%) | 0.746 |
|  | wo/Rec | 9 (31.0%) | 6 (37.5%) |  |
| **Recipient age** (mean (±SD)) |  | 53.14 (±13.20) | 54.25 (±10.43) | 0.773 |
| **Recipient gender** | Female | 9 (31.0%) | 4 (25.0%) | 0.743 |
|  | Male | 20 (69.0%) | 12 (75.0%) |  |
| **Recipient race** | Black | 22 (75.9%) | 13 (81.2%) | 1 |
|  | White | 5 (17.2%) | 3 (18.8%) |  |
|  | Hispanic | 1 (3.4%) | 0 (0.0%) |  |
|  | Other | 1 (3.4%) | 0 (0.0%) |  |
| **Etiology of ESRD** | DM | 5 (17.2%) | 2 (12.5%) | 0.898 |
|  | DM/HTN | 7 (24.1%) | 6 (37.5%) |  |
|  | FSGS | 3 (10.3%) | 2 (12.5%) |  |
|  | HTN | 11 (37.9%) | 5 (31.2%) |  |
|  | Other | 3 (10.3%) | 1 (6.2%) |  |
| **Donor type** | DCD | 3 (6.67%) | 5 (11.11%) | 0.11 |
|  | non-DCD | 26 (57.78%) | 11 (24.44%) |  |
| **KDPI** (% , mean (±SD)) |  | 48.45 (±23.58) | 61.56 (±22.75) | 0.078 |
| **Cause of Death** | Anoxia | 10 (34.5%) | 8(50.0%) | 0.098 |
|  | Head Trauma | 9 (31.0%) | 1(6.2%) |  |
|  | Stroke | 7 (24.1%) | 7(43.8%) |  |
|  | other | 3 (10.3%) | 0(0.0%) |  |
| **Donor age** (years, mean (±SD)) |  | 36.59 (±15.47) | 39.62 (±14.90) |  |
| **Donor gender** | Female | 14 (48.3%) | 5 (31.2%) | 0.351 |
|  | Male | 15 (51.7%) | 11 (68.8%) |  |
| **Donor race** | Asian | 0 (0.0%) | 1 (6.2%) | 0.097 |
|  | Black | 16 (55.2%) | 4 (25.0%) |  |
|  | White | 13 (47.8%) | 11 (68.8%) |  |
| **Donor HCV Ab** (%) | Negative | 28 (96.6%) | 14 (87.5%) | 0.285 |
| **Donor terminal creatinine** mg/dL (mean (±SD)) |  | 1.53 (±0.98) | 1.69 (±1.15) | 0.31 |
| **Cold ischemia time** in min (mean (±SD)) |  | 1071.41 (±319.64) | 1135.19 (±214.30) | 0.48 |
| **eGFR** (mean (±SD))<br>measured in mL/min/1.73m<sup>2</sup> | **1 week** | 51.15 (±29.79) | 12.48 (±5.06) | <0.001 |
|  | **1 month** | 64.22 (±22.22) | 38.90 (±16.53) | <0.001 |
|  | **3-month** | 61.05 (±23.12) | 44.13 (±19.58) | 0.017 |
|  | **6-month** | 60.06 (±19.12) | 44.38 (±19.95) | 0.013 |
|  | **12-month** | 57.83 (±20.93) | 45.79 (±17.43) | 0.063 |
| | **24-month** | 60.25 ($\pm 24.59$) | 48.98 ($\pm 21.67$) | 0.144 |

**Abbreviations:** PostKT-non-DGF: post kidney transplantation without delayed graft function; PostKT-DGF: post kidney transplantation with delayed graft function; post kidney transplantation without delayed graft function; w/Rec: with functional recovery; wo/Rev: without functional recovery; DCD: donation after circulatory death; ESRD: end stage renal disease; KDPI: Kidney Donor Profile Index; DM/HTN: Diabetes Mellitus/Hypertension; DM: Diabetes Mellitus; HTN: Hypertension; FSGS: Focal segmental glomerulosclerosis; A: Asian; AA: African American; C: Caucasian; H: Hispanic; O: Other; eGFR: estimated glomerular filtration rate measured in mL/min/1.73m<sup>2</sup>.

### Functional outcomes and histological analysis of kidney transplants

Among the postKT-AKI cases, three patients (AKI465, AKI560, and AKI573) showed functional recovery (postKT-AKIw/Rec), with follow-up eGFR > 50 mL/min/1.73m² (range: 55– 80) at 24 months and a sustained upward trend posttransplant (**Table 1**), as previously defined (*23*). Notably, this included a kidney with a high KDPI of 76% (AKI573) and another with a donor terminal creatinine of 5.7 mg/dL (AKI560). In contrast, four patients (AKI306, AKI505, AKI538, and AKI542) experienced persistent dysfunction (postKT-AKIwo/Rec), with eGFR ≤30 mL/min/1.73m². One patient (AKI505) experienced allograft loss and returned to chronic hemodialysis 22 weeks post-KT. The remaining three maintained eGFR levels between 19–23 mL/min/1.73m² at 24 months post-KT.

Sequential biopsy analysis in postKT-AKIwo/Rec patients revealed progressive tubulointerstitial damage. Patient AKI505 developed severe interstitial fibrosis and tubular atrophy (IFTA) without evidence of rejection. AKI306 progressed from no fibrosis (ci=0, ct=0) at baseline to moderate interstitial IFTA (ci=2, ct=2) at 4 months post-KT. AKI538 developed borderline T cell-mediated rejection (TCMR) at 15 weeks post-KT, which responded to steroid therapy. AKI542 exhibited moderate IFTA and early transplant glomerulopathy on electron microscopy at 6 months. Characteristics and histological features of nKTx (collected > 24 months post-KT) are summarized in **Table 2**. This well-characterized cohort enabled direct comparisons of early injury responses across clinically relevant transplant conditions.

### Single-nucleus transcriptomic profile of renal cell types in kidney biopsies

We performed snRNA-seq on kidney biopsies and integrated transcriptomic data after batch correction. Following rigorous quality control (QC), 38,534 high-quality nuclei were retained (**Fig. S1**, **Tables S1**, **S2**). A total of 19 major cell clusters were identified, representing proximal tubules (PT1–4), distal tubules (DT1–3), collecting duct principal (CDP1 & 2), collecting duct intercalated (CDI1 & 2), endothelial cells (EC1–4), podocytes (POD), fibroblasts (Fib), a transitioning cell cluster (TC), and immune (IMM) cells (**Fig. 1B, Fig. S2, Table S3**). Marker genes for each cell cluster were identified by comparing cells within a specific cluster against all remaining cells (|fold change| (FC) > 1.5; adjusted (adj.) p-value < 0.01) using the CellMarker database (*24, 25*) (**Fig. 1C**, **Table S4**). The proportion of cells in each cluster and within each group is depicted in **Fig. 1D**. Clusters containing fewer than five marker genes were considered over-clustering artifacts and excluded from downstream analyses.

### Proximal tubule cell clusters and injury response

PT cells were most abundant in all conditions. Four proximal tubule PT subclusters (PT1 to PT4) comprising 11,712 cells were identified (**Fig. 2A**). PT1 was enriched in NNK (19%) but reduced in nKTx (9%) and remained similarly low in both postKT-AKIw/Rec and postKT-AKIwo/Rec (6.3% vs. 6.5%) (**Fig. 1D**). PT2 was more prevalent in postKT-AKIw/Rec than in postKT-AKIwo/Rec (8.4% vs. 5.4%), whereas PT3 expanded from 5.1% in postKT-AKIwo/Rec to 10.7% postKT-AKIw/Rec, despite comparable proportions in NNK and nKTx. PT4 showed a stepwise increase from 3% in NNK to 10.8% in postKT-AKIwo/Rec. Additionally, a fifth population, termed transitional cells (TC), expressed markers of both tubular and fibroblast. TC were markedly enriched in postKT-AKIwo/Rec (7.1%) compared to postKT-AKIw/Rec (4.5%), nKTx (2.7%), and NNK (1.4%) (**Fig. 1D**). Gene set enrichment analysis of this cluster revealed activation of pathways involved in cell-cell adhesion, cytoskeletal remodeling, tube morphogenesis, and nephrogenesis (**Fig. S3**), consistent with a transitional or injury-responsive state.

**Fig 2.**
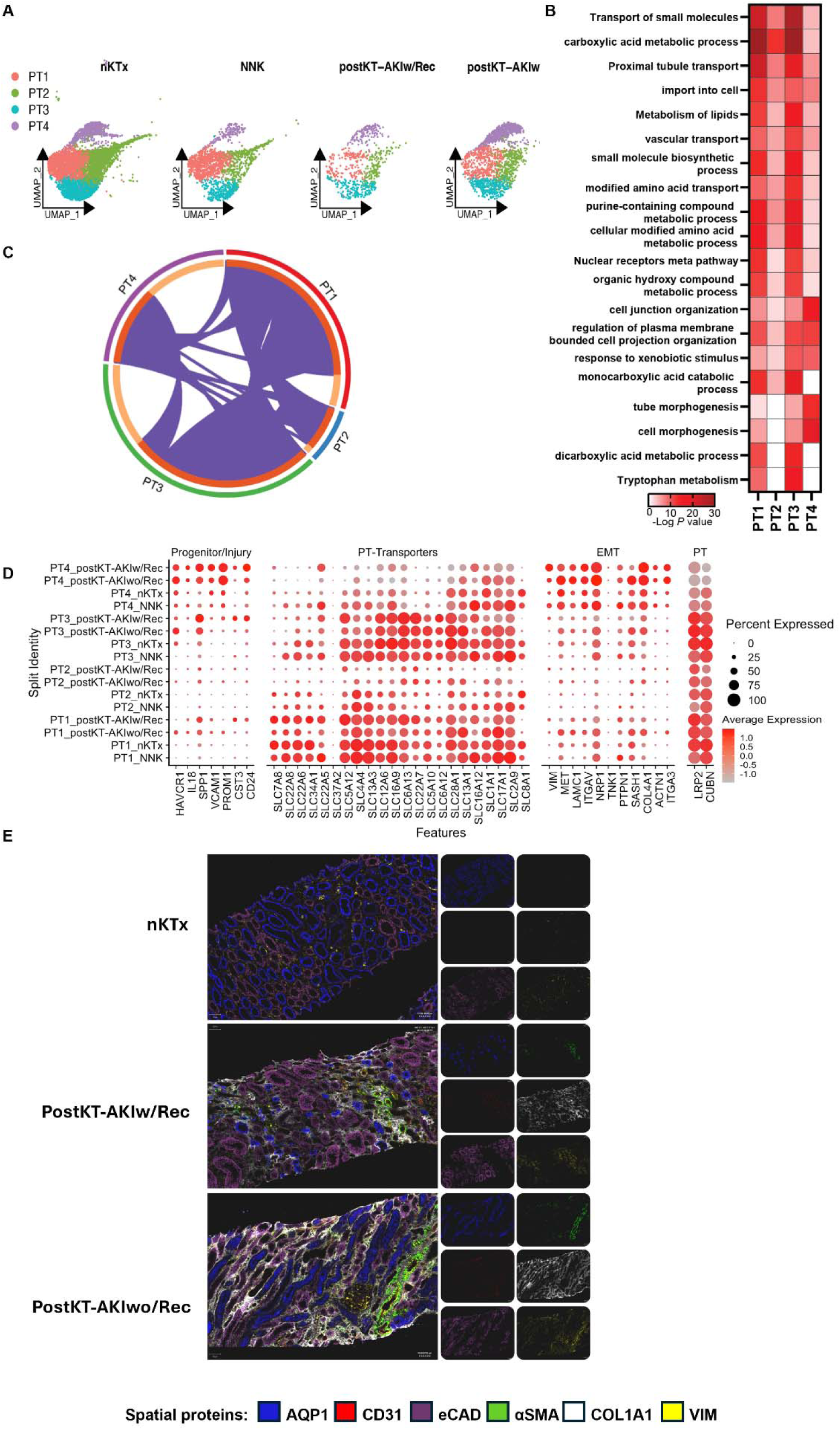
Functional properties of PT clusters. (**A**) Four PT subclusters identified across different conditions. (**B**) Biological functions and pathways enriched for the PT marker genes associated with each subcluster (PT1-4). (**C**) Circos plot illustrating the degree of overlap between marker related genes between the clusters indicates that PT1 and PT3 are enriched for functions minimally shared with the altered PT cells of the PT4 subcluster. (**D**) Dot plot demonstrates the expression pattern of progenitors, transporters, EMT, and normal proximal tubule genes in PT clusters. (**E**) Imaging mass cytometry data from nKTx, postKT-AKIw/Rec and postKT-AKIwo/Rec samples for tubule marker (AQP1, eCAD), endothelial cells (CD31), ECM (COL1A1, αSMA) and EMT (VIM) for structure assessment of the kidney biopsies.

Marker gene ontology (GO) analysis (adj. p < 0.01, |log FC| > 0.5; **Table S5**) revealed distinct biological processes across PT subclusters, despite a shared core of proximal tubule-associated genes confirming their common lineage. PT1 was enriched for metabolic and solute transport pathways, consistent with healthy proximal tubule function and minimal stress signaling. PT2 retained core PT functions but also showed features of early adaptation, including enrichment of small molecule transport and carboxylic acid metabolism pathways (**Fig. 2B**). Of note, PT2 showed increased expression of *SLC8A1* (*NCX1*), a calcium exporter implicated in stabilizing E-cadherin at cell junctions, suggesting preserved epithelial integrity despite early injury signals (*26*). PT3 displayed reduced expression of transport and lipid metabolism genes, indicating a transitional state toward dysfunction, with an intermediate phenotype also reflected in PT2 and PT4 clusters (**Fig. 2C**). PT4 was enriched for developmental, cytoskeletal, and receptor tyrosine kinase signaling, consistent with dedifferentiation and stress-activated transcriptional programs. PT4 showed the most significant transcriptional reprogramming, marked by near-complete loss of solute transport, metabolic, and energy-related pathways (**Fig. 2D**). Together, these data position PT1 cell cluster as a homeostatic reference, PT2 as an adaptive/stable phenotype, PT3 as a transitional state, and PT4 as a potentially maladaptive or pre-fibrotic state.

IMC analysis corroborates transcriptional findings of tubular injury in post-KT AKI samples (**Fig. 2E**). We then evaluated the observed transcriptional changes in PTs at the protein level using IMC in five slides from same tissue used for snRNA-seq (1 nKTx, 1 postKT-AKI w/Rec, and 3 postKT-AKI wwo/Rec). Compared to intact architecture in nKTx, postKT-AKI samples showed pronounced disruption of epithelial integrity, evidenced by reduced E-cadherin (eCAD) and AQP1 staining, hallmarks of PT identity. Progressive upregulation of α-smooth muscle actin (αSMA), vimentin (VIM), and collagen type I (COL1A1) was observed, particularly in structurally compromised regions which was increased in postKT-AKIwo/Rec as quantified and discussed in the **Supplementary Information**. These spatial patterns suggest ongoing fibrotic remodeling and epithelial-to-mesenchymal transition (EMT). Co-localization of mesenchymal and fibrotic markers with eCAD loss supports the emergence of TC cells and PT4-like phenotypes, reinforcing the transcriptional evidence for maladaptive epithelial states in grafts that fail to recover.

### PT subcluster transcriptional profiles across conditions

#### nKTx vs. NNK: Adaptation of transplanted kidneys to a new microenvironment

The cellular response of PT cells to transplant-related stress determines whether epithelial repair proceeds toward functional recovery or culminates in maladaptive remodeling and chronic graft injury. To define programs associated with structurally stable allografts, we compared differentially expressed genes (DEGs) across PT subclusters between nKTx and NNK (**Fig. 3A, Table S6**). While all PT clusters retained core proximal tubule identity, each exhibited unique molecular adaptations. In nKTx, transcriptional changes were consistent with functional adaptation to the allogeneic environment. PT1 showed upregulation of vesicle-mediated transport and AMPK signaling, suggesting enhanced intracellular trafficking and metabolic homeostasis (**Fig. 3A**). PT2 was enriched for Notch3 signaling and apical–basal polarity maintenance, reinforcing epithelial organization and function. In PT3, the activation of the VEGFA-VEGFR2 signaling implied endothelial crosstalk and support for vascular remodeling in the kidney graft. In contrast, PT4 exhibited downregulation of solute transport, kidney development, and carboxylic acid metabolism pathways, indicating reduced metabolic activity. While PT1-PT3 maintained features of native tubules, the transcriptional profile of PT4 may reflect subtle injury or stress, potentially driven by persistent subclinical immune alloreactivity, as we previously reported (*27*).

**Fig 3.**
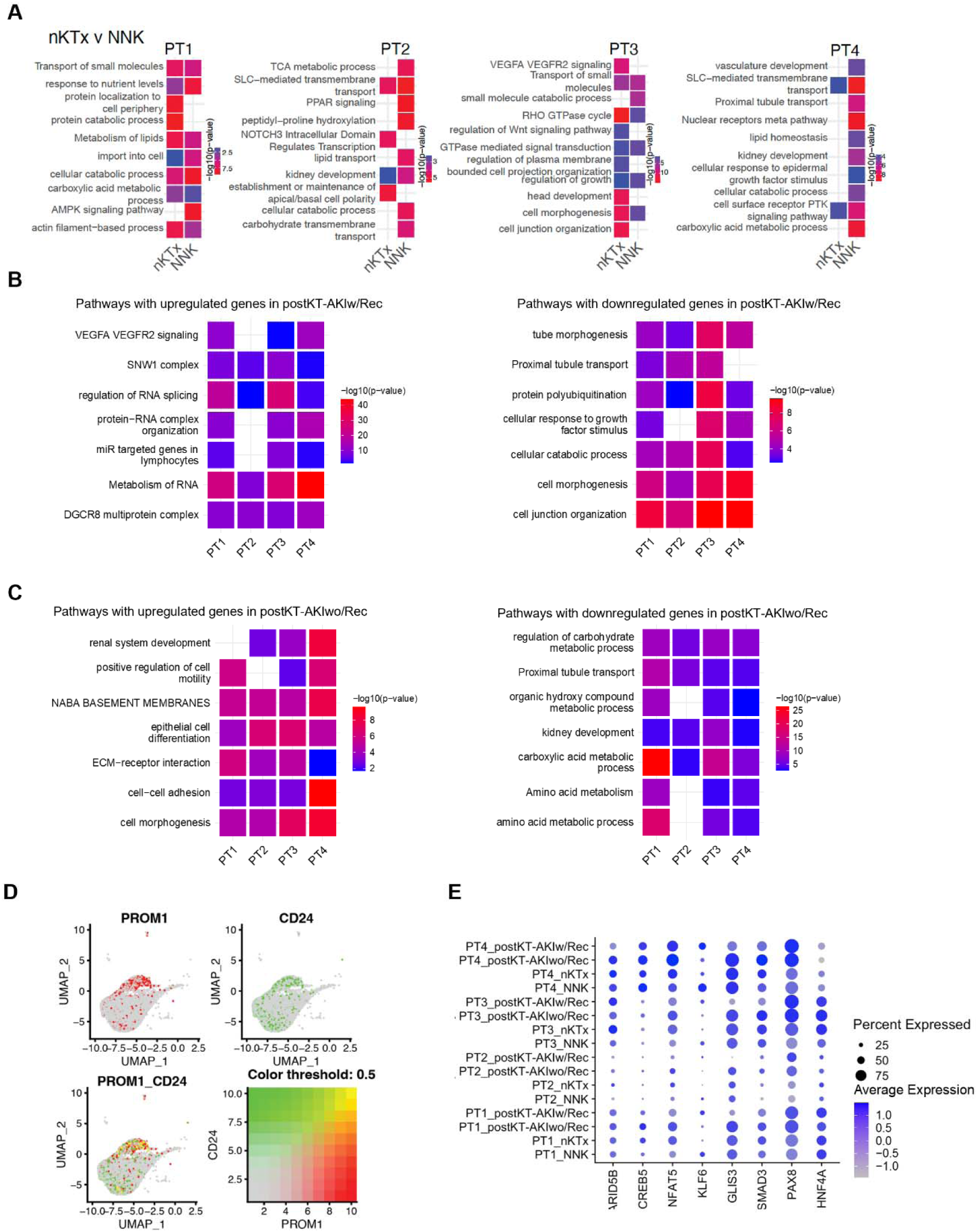
Differential properties of PT cell clusters. The DEGs were used to determine the enrichment of pathways in the (**A**) NNK v nKTx, (**B**) postKT-AKIw/Rec v nKTx and (**C**) postK-AKIwo/Rec v nKTx. The pathways enriched are based on overexpressed genes in that condition. Metascape was used to identify the enriched pathways. (**D**) Expression and co-expression analysis of *PROM1* and *CD24* in PT subclusters in an integrated UMAP. The left panel shows *PROM1* expression, which is highest in PT4. The middle UMAP panel illustrates *CD24* expression, with PT4 showing the highest levels, followed by PT3. The bottom panel demonstrates the co-expression of *PROM1* and *CD24*, which is predominantly increased in the PT4 subcluster. (**E**) Dot plot representation for the expression of transcription factor in the different condition.

#### postKT-AKIw/Rec vs. nKTx: Partial recovery with persistent cellular stress

In postKT-AKIw/Rec grafts, PT subclusters partially retained the adaptive transcriptional signatures observed in nKTx but also demonstrated stress-related responses. PT1 and PT2 were enriched for pathways related to RNA metabolism, splicing regulation, and protein–RNA complex assembly, reflecting roles in post-injury homeostasis and adaptive recovery (**Fig. 3B, Table S7**). PT3 was enriched for translational control, VEGFA–VEGFR2 signaling, and miRNA-regulated gene sets, indicating a transitional state between adaptation and stress.

PT4 in this group showed broad transcriptional suppression across pathways governing metabolic activity, protein localization, translation, and transmembrane transport, particularly of SLC-family transporters essential for proximal tubule function. These patterns suggest persistent dedifferentiation and incomplete repair. Despite apparent improvement in renal function at the graft organ level, the transcriptional profile of PT4 reflected a maladaptive state, highlighting the disconnect between functional metrics and cellular stress resolution. This disconnect supports the development of molecular biomarkers or transcriptomic signatures to better stratify patient risk for chronic graft dysfunction or fibrosis, especially when standard clinical indicators (e.g., serum creatinine, eGFR) fail to detect subclinical injury. Moreover, identifying patients with maladaptive PT signatures (like PT4) despite stable function could allow for targeted immunomodulatory or antifibrotic therapies before overt decline occurs.

#### postKT-AKIwo/Rec vs. nKTx: Progressive dysfunction and fibrotic remodeling

In contrast to recovering grafts, PT cells from postKT-AKIwo/Rec samples demonstrated transcriptional signatures of ongoing dysfunction across all PT subclusters. Upregulated genes in PT1, PT3, and PT4 clusters were enriched for pathways linked to cell motility and adhesion. Concurrently, all clusters showed downregulation of pathways related to carboxylic acid metabolism, solute transport, and nephron development, indicating broad suppression of physiological function (**Fig. 3C).** This population uniquely upregulated pathways related to extracellular matrix (ECM) remodeling, epithelial-to-mesenchymal transition (EMT), and fibrosis, including NABA basement membrane remodeling, consistent with a fibrogenic shift.

Canonical PT markers (e.g., *CUBN, LRP2*) and essential transporters were markedly reduced, further supporting a dedifferentiated, injury-associated phenotype. Interestingly, PT4 cell cluster also co-expressed *PROM1* (*CD133*) and *CD24* (**Fig. 3D**), markers of renal progenitor-like cells, suggesting that even within a maladaptive environment, a regenerative subpopulation may persist. These cells also upregulated transcription factors (TFs) associated with injury and repair (*ARID5B, CREB5, NFAT5, PAX8, KLF6*) while downregulating *HNF4A*, a master regulator of proximal tubule identity and repair (*28*) (**Fig. 3E**).

The expansion of PT4 in postKT-AKIwo/Rec, coupled with enriched profibrotic and dedifferentiated signatures, indicates a failure to resolve injury. This divergence from the postKT-AKIw/Rec profile suggests the presence of a critical transcriptional checkpoint, where PTs either re-establish homeostasis or transition toward chronic injury and fibrosis.

### Distinct fibroblast clusters and divergent trajectories drive fibrosis and repair in post-transplant acute kidney injury

snRNA-seq revealed a transcriptionally diverse fibroblast population (Fib; n=1,658 nuclei), subclustered into mural cells, mesangial cells (MES), myocytes (MYO), and seven fibroblast states: stromal progenitors (StrProg), quiescent fibroblasts (qFIB), medullary fibroblasts (mFIB), two activated interstitial fibroblast subtypes (aFIB1 and aFIB2), myofibroblasts (MYOF), and proliferating fibroblasts (pFIB) (**Fig. 4A-B**). These clusters represent a continuum from quiescent to fibrogenic states. Differential expression and pathway analysis identified pFIB as a highly mitotic population, enriched in cell cycle genes, while MYOF and aFIB2 expressed gene sets associated with ECM remodeling and fibrosis (**Fig. 4CD**). In contrast, qFIBs (predominant in nKTx) lacked ECM-related gene expression, consistent with a homeostatic role (“resting” fibroblasts).

**Fig 4.**
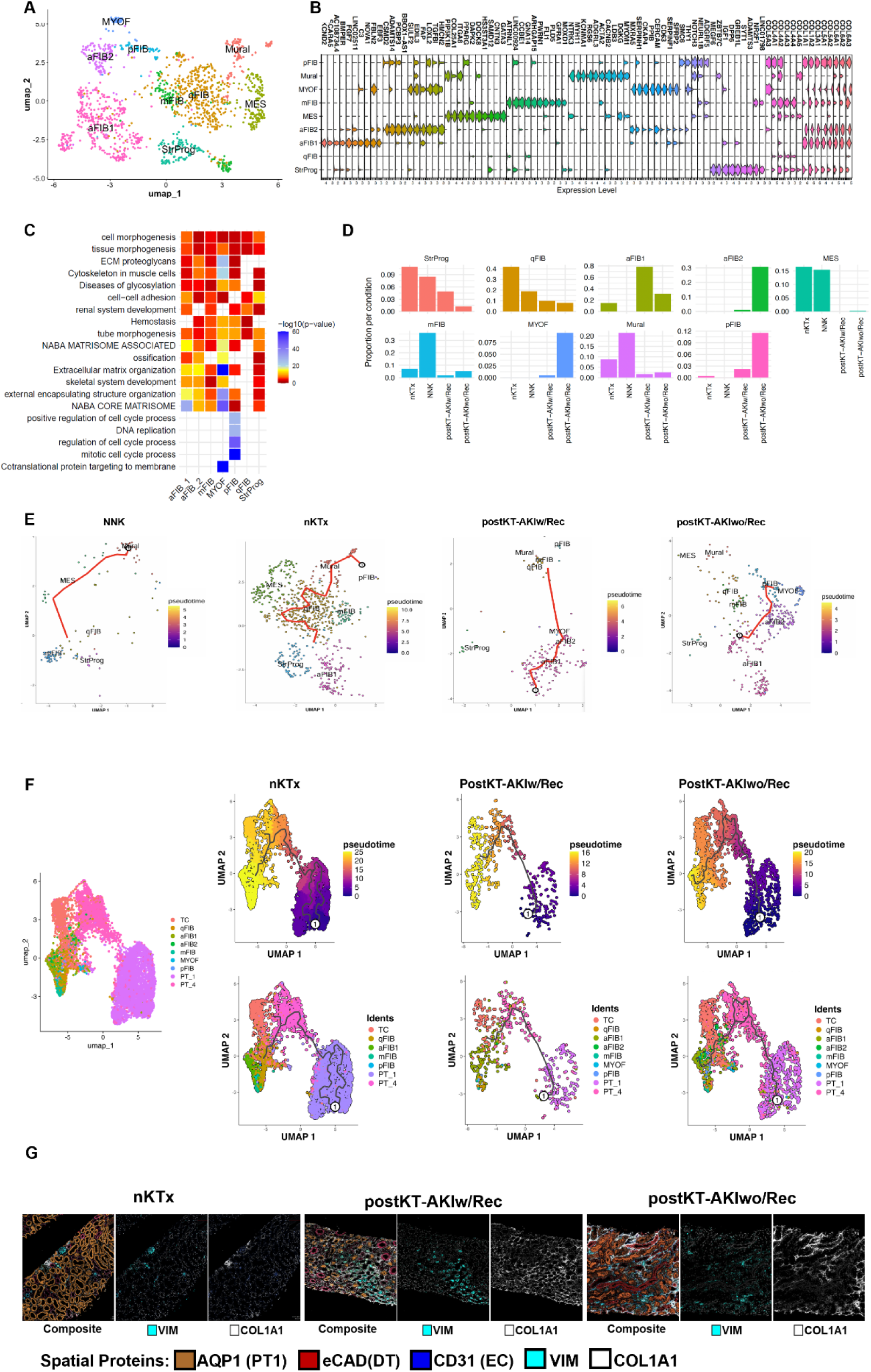
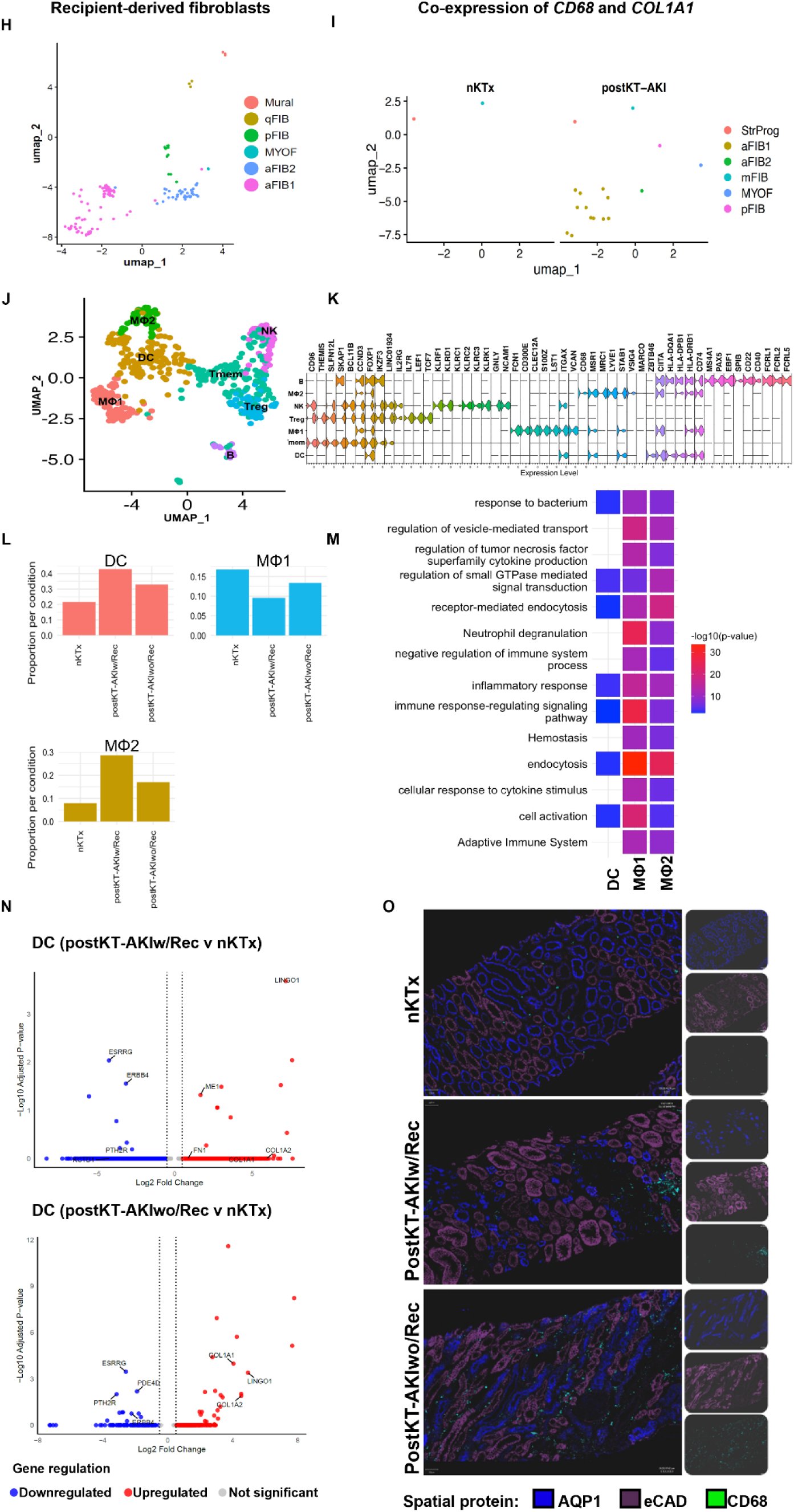
Sub-clustering of Fibroblast and trajectory analysis. (**A**) Integrated UMAP of nine Fib sub-clusters. (**B**) Violin plots for the marker genes used for classification (**C**) Biological pathways enriched for each subcluster marker genes (**D**) Proportion of nuclei in each sub-cluster across different conditions. (**E**) Single cell pseudotime trajectory revealed distinct lineage progressions among Fib subpopulations. (**F**) Trajectories along pseudotime for the nKTX, postKT-AKIw/Rec and postKT-AKIwo/Rec depicting the transition of PT4 cells to MYOF and activated Fib (right). Solid black line depicts the expression curves for each branch over pseudotime. An encircled 1 indicates the origin of trajectory. (**G**) IMC images with VIM and COL1A1supporting the transition of injured PT to Fib-like cells. (**H**) UMAP of recipient derived Fib nuclei in sex-mismatched transplants (**I**) Co-expression of *CD68* (macrophage) and *COL1A1* (fibroblast) in Fib (**J**) Integrated UMAP visualization of immune cell sub-clusters. (**K**) Violin plots for the marker genes used to classify immune sub-clusters. (**L**) Proportion of nuclei in MΦ and DC subcluster across different conditions. (**M**) Biological pathways enriched for the MΦ and DC subcluster marker genes (**N**) Volcano plot for DEGs in DC in postKT-AKIw/Rec and postKT-AKIwo/Rec compared to nKTx. (**O**) IMC confirmed CD68+ macrophages in matched kidney biopsies.

Fib subtype distribution varies markedly across transplant conditions. aFIB2, MYOF, and pFIB were exclusive to postKT-AKIwo/Rec, whereas qFIB and StrProg were enriched in nKTx (**Fig. 4D**). Trajectory analysis revealed divergent fibroblast lineages between PostKT-AKIw/Rec and wo/Rec. In nKTx, StrProg transitioned into aFIB1 but did not progress toward profibrotic states, suggesting a restrained, non-pathological activation (**Fig. 4E**). Mural cells contributed to both qFIB and mFIB lineages, supporting their structural role in tissue maintenance. In postKT-AKIwo/Rec, continuous progression from aFIB1 to aFIB2 and MYOF was observed, defining a fibrogenic trajectory. This transition was markedly attenuated in postKT-AKIw/Rec, where aFIB2 and MYOF states were largely absent, supporting a model of limited fibroblast activation associated with successful recovery.

To characterize injury-associated cell state transitions, we first performed integrated clustering of PT and Fib populations across conditions (**Fig. S4**). Subpopulations with the greatest spatial and transcriptional relevance to injury and repair were selected for focused pseudotime analysis. An integrated UMAP and pseudotime trajectory analysis was then conducted on these selected epithelial and fibroblast clusters across postKT-AKIw/Rec, postKT-AKIwo/Rec, and nKTx samples (**Fig. 4F**). Cells were annotated into nine subpopulations including TC, qFIB, aFIB1, aFIB2, mFIB, MYOF, pFIB, and PT clusters PT1 and PT4. In nKTx, pseudotime analysis revealed a branched trajectory terminating in PT1 and qFIB, consistent with epithelial and stromal homeostasis. PostKT-AKIw/Rec displayed a moderately extended trajectory, with expansion of TC and aFIB1 but limited progression into aFib2 or MYOF, consistent with partial remodeling and controlled fibroblast activation. In contrast, postKT-AKIwo/Rec exhibited a linear, unidirectional trajectory culminating in PT4 and fibrogenic fibroblast states (aFIB2, MYOF, pFIB), reflecting maladaptive injury and persistent fibrosis. The integrated UMAP embedding showed spatial segregation of regenerative versus fibrotic endpoints, with PT4 closely aligned with aFIB2 and MYOF, supporting the hypothesis of epithelial-stromal crosstalk in driving fibrotic remodeling in non-recovery grafts.

IMC confirmed spatial localization of fibrosis and mesenchymal activation across conditions. In nKTx, AQP1 PT cells retained epithelial organization with minimal COL1A1 or VIM expression, consistent with a homeostatic epithelial and stromal environment. In postKT-AKIw/Rec, moderate peritubular VIM and focal COL1A1 expression were observed, reflecting localized injury and early remodeling, but limited fibrotic progression. In contrast, postKT-AKIwo/Rec demonstrated diffuse COL1A1 deposition and robust VIM expression in AQP1 tubules, particularly at the fibrotic interface, consistent with partial EMT and active fibrotic remodeling (**Fig. 4G**).

To investigate fibroblast origin, we analyzed sex-mismatched donor-recipient kidneys using *XIST* (X chromosome) and *KDM5D* (Y chromosome) expression. Recipient-derived fibroblasts were enriched in the aFIB1 cluster and co-expressed *CD68* and *COL1A1* (**Fig. 4H–I**), consistent with macrophage (MΦ)-to-MYOF transition (MMT), as previously described (*29*). These MMTs may contribute to Fib accumulation in postKT-AKI, where Fib comprised 8% of cells, compared to 3.3% in nKTx and 2.5 in NNK.

DCs and two MΦ subsets (MΦ1, MΦ2) were identified (**Fig. 4J–M**). DCs were most abundant in postKT-AKIw/Rec and expressed reparative genes (*ARHGAP22, CSF1R, HLA-DQA1)* (**Fig. 4N**). In contrast, DCs in postKT-AKIwo/Rec exhibited upregulation of fibrosis-associated genes (*COL1A1, COL1A2*) and downregulation of metabolic and reparative pathways (*PDE4D, PTH2R, ESRRG*), indicating a profibrotic shift (*30–35*).

MΦ1s, marked by *FCN1, CD300E*, and *CLEC12A,* were enriched in postKT-AKIwo/Rec (13.4%) and associated with chronic inflammation (*36*). MΦ2s, expressing *CD163, IL10RA, TGFBR1/2,* and *MS4A7*, were most abundant in postKT-AKIw/Rec (28.6%) and aligned with tissue repair (*37, 38*). IMC confirmed spatial colocalization of COL1A1 fibrotic zones with VIM PTs and immune infiltrates, suggesting immune-stromal interaction drives maladaptive remodeling in postKT-AKIwo/Rec (Fig. **4O**). The exclusive emergence of aFIB2, MYOF, and MTM populations in postKT-AKIwo/Rec defines a transcriptional and spatial program of persistent injury and fibrotic remodeling.

### Adaptive *versus* maladaptive pathways in the proximal tubular cells associate with divergent outcomes in kidney grafts

Our dataset includes KTRs who experienced either postKT-AKIw/Rec or postKT-AKIwo/Rec, enabling direct comparison of transcriptional profiles associated with favorable vs. maladaptive outcomes. We profiled 577 PT cells from postKT-AKIw/Rec and 2,097 from postKT-AKIwo/Rec samples, encompassing PT1-PT4 clusters.

DEG analysis comparing all PT clusters between the two conditions (FC > |0.5|, adj. p < 0.05) revealed distinct transcriptional programs (**Fig. 5A, Table S10**). In PT2 cells from postKT-AKIwo/Rec, upregulated genes were enriched for tube morphogenesis, Wnt signaling regulation, cell projection organization, and cell morphogenesis, suggesting aberrant activation of developmental and morphogenetic pathways (**Fig. 5B**). In contrast, PT3 and PT4 cells in postKT-AKIw/Rec were enriched for pathways associated with protein synthesis, biological oxidation, and amino acid metabolism, consistent with adaptive recovery and energy reprogramming. Shared pathway enrichments between PT3 and PT4 in postKT-AKIw/Rec suggested functional similarities within the recovering nephron (**Fig. 5B**).

**Fig 5.**
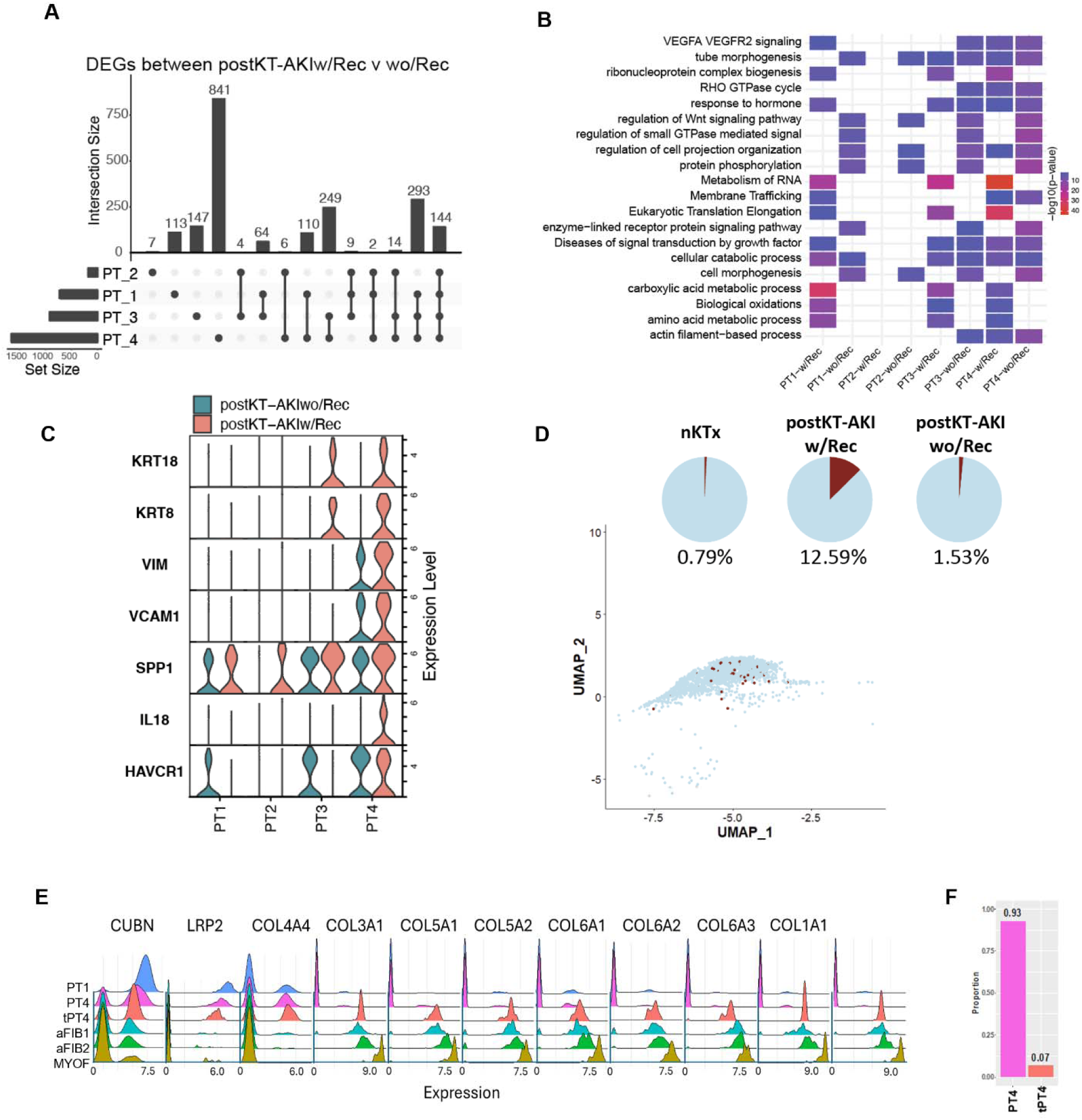
Transcriptional heterogeneity in PT cluster between postKT-AKIw/Rec and postKT-AKIwo/Rec. **(A**) UpSet plot visualization for DEGs in PT clusters (PT1-PT4) between postKT-AKIw/Rec and postKT-AKIwo/Rec, the intersection size depicts the number of genes commonly differentially expressed across the dataset. The filled dots and connecting lines indicate th DEGs shared by those clusters. (**B**) Biological functions and pathways enriched for DEGs in each PT cluster. The pathways enriched are based on the overexpressed genes in that condition. (**C**) Violin plot for injury markers (**D**) UMAP for the co-expression of STC markers (*PROM1^+^ CD24^+^ VIM^+^ PAX2^+^ HAVCR1^+^ ALDH1A1^+^),* each brown dot represents a nucleus co-expressing the markers. The pie chart depicts the proportion of these cells in different conditions. (**E**) Ridge pot to visualize the distribution of genes across PT and Fib clusters. Higher peaks in similar location indicate similar expression pattern of the gene in those clusters. The tPT4 is a subset of transitioning PT4 cells characterized by loss of PT markers (*CUBN*, *LRP2*) and gain of fibroblast markers *COL4A4, COL3A1, COL5A1, COL5A2, COL6A1, COL6A2, COL6A3, COL1A1.* (**F**) The bar graph on the right depicts the proportion of tPT4 in postKT-AKIwo/Rec.

In postKT-AKIwo/Rec, all PT clusters showed broad enrichment for phosphorylation signaling, Wnt pathway regulation, small GTPase-mediated signal transduction, and enzyme-linked receptor signaling, consistent with sustained activation and impaired resolution of injury. These findings suggest that cellular plasticity in postKT-AKIwo/Rec is skewed toward maladaptive remodeling rather than repair.

We next examined injury markers’ expression across conditions. *HAVCR1* (KIM-1) expression was higher in PT4 cells from postKT-AKIwo/Rec, while *VIM, VCAM1, IL18*, and *SPP1* were more highly expressed in PT4 cells from postKT-AKIw/Rec (**Fig. 5C**). Co-expression of *HAVCR1* and *VIM* in PTs has been linked to epithelial proliferation during repair (*39, 40*), while *VCAM1* facilitates immune cell recruitment and *IL18* serves as an inflammatory mediator. Notably, *KRT8* and *KRT18* were also upregulated in PT4 cells from postKT-AKIw/Rec, suggesting engagement of regenerative programs (*40*). Based on this transcriptional signature, we defined PT4 cells in postKT-AKIw/Rec as injured proinflammatory PT cells, which represent a transitional state engaged in epithelial repair.

In postKT-AKIw/Rec, the PT4 cluster was enriched for co-expression of *PROM1, CD24, VIM, PAX2, HAVCR1,* and *ALDH1A1*, a gene signature consistent with scattered tubular cells (STCs) (**Fig. 5D**). STCs are a progenitor-like population with established proliferative and regenerative potential (*41–44*). Quantification of this subset revealed a 12.6% STC composition in postKT-AKIw/Rec, compared to only 1.5% in postKT-AKIwo/Rec and 0.8% in nKTx, highlighting a markedly enhanced regenerative response in recovering grafts.

In contrast, a subset of PT4 cells in postKT-AKIwo/Rec exhibited transitional features characterized by loss of PT identity genes (*CUBN, LRP2*) and upregulation of fibroblast-associated genes, including *COL1A1, COL5A1/2, COL6A1-3, COL4A4* (**Fig. 5E**). These cells transcriptionally overlapped with aFIB, aFIB2, and MYOF clusters, indicating a shift toward a mesenchymal-like, profibrotic phenotype. We designated this population as transitional PT4 (tPT4) cells. tPT4 cells accounted for 7% of the injured PT compartment and were exclusively observed in postKT-AKIwo/Rec samples (**Fig. 5F**). Together, these data highlight a bifurcation in the fate of injured PT cells showing that in recovery, PT4 cells engage a proinflammatory but regenerative STC program, while in non-recovery, a distinct subset transitions toward fibrotic reprogramming, reinforcing maladaptive tissue remodeling.

### Cell–Cell crosstalk between tubular, fibroblast, and immune compartments in post-transplant AKI

The pairwise chord plots illustrate ligand-receptor (LR) interactions among PT, Fib, and immune cells (**Fig. 6, Table S11, Fig. S5**). In postKT-AKI w/Rec, the strongest LR interactions were observed between aFIB1 and PT4, particularly the PAM-FAP interaction, which may drive Fib activation and fibrosis. However, this interaction was more pronounced in postKT-AKI wo/Rec, involving aFIB1, aFIB2, and MYOF with PT4. Also, in postKT-AKI wo/Rec, Fib subclusters (aFIB1, aFIB2, MYOF, pFIB) exhibited PAM-FAP interactions among themselves, potentially sustaining FAP activation and contributing to maladaptive repair post-AKI (*12*). The COL18A1-ITGA5 interaction (*45*), was observed between PT4 and aFIB1 in postKT-AKI w/Rec, whereas in postKT-AKI wo/Rec, it was more broadly distributed between PT and aFIB2/MYOF, suggesting a more persistent fibrotic signaling in the latter condition.

**Fig 6.**
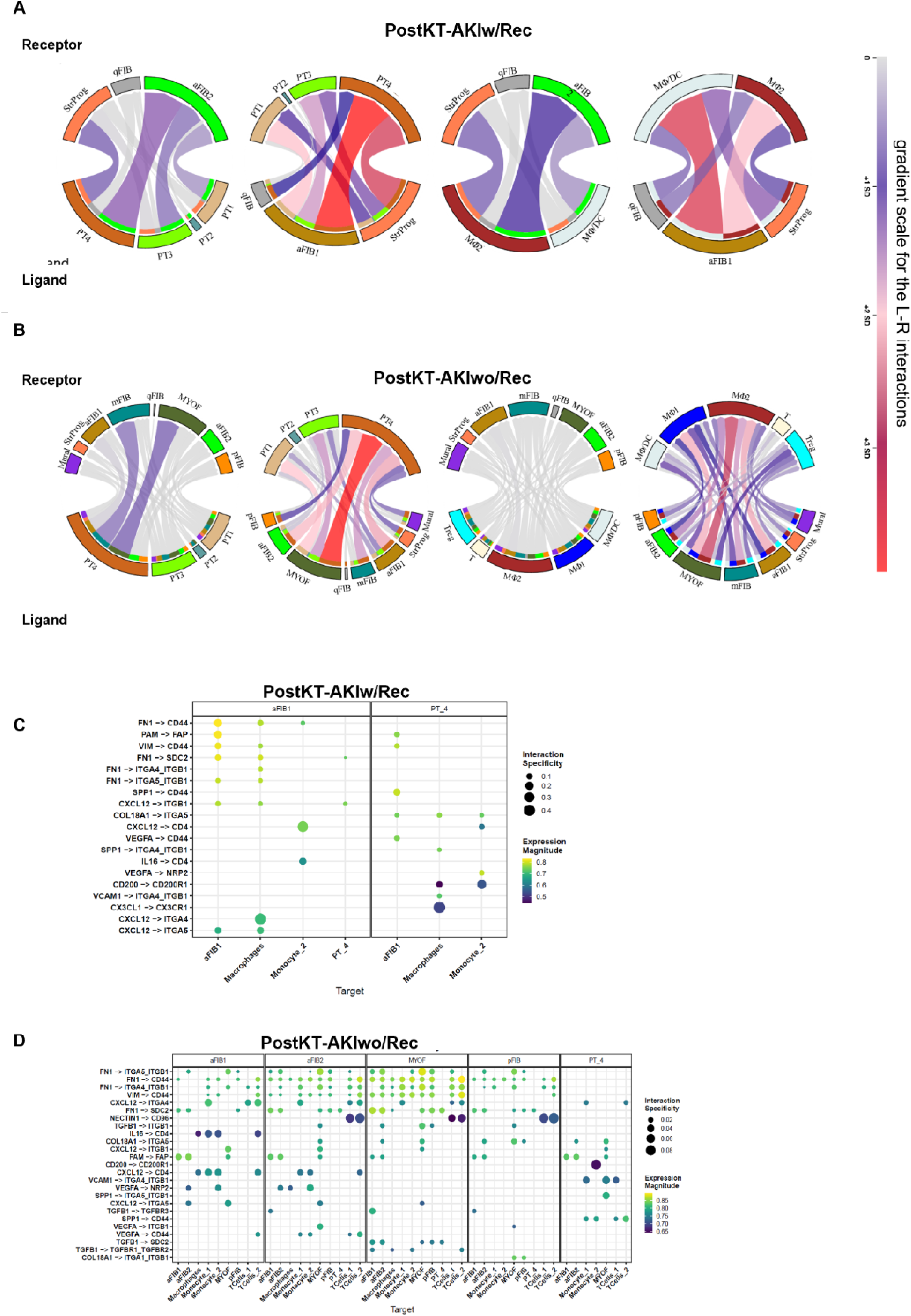
Ligand receptor interactions. Chord plots for LR interactions between PT, Fib and Immune cells (A) in postKT-AKIw/Rec and (B) postKT-AKIwo/Rec. Chords between ligands and receptors are colored according to the interaction strength, based on standard deviation (SD) from the mean. Bubble plot showing interaction data related to different LR pairs (C) in postKT-AKIw/Rec and (D) postKT-AKIwo/Rec.

In postKT-AKI w/Rec, significant interactions between fibroblast and immune sub-clusters were observed, particularly between aFIB1 and macrophages (MΦ/DC, MΦ2). The LR pair CXCL12-ITGA4, identified between aFIB1 and MΦ1, is known to recruit macrophages to sites of inflammation and regions with high ECM deposition (*46*). Notably, the CD200-CD200R1 and CX3CL1-CX3CR1 interactions between the PT4 cluster and the MΦ1 sub-cluster play a crucial role in dampening inflammation by suppressing pro-inflammatory cytokine expression (*47, 48*). LR interactions involving adhesion molecules and integrins, such as VIM-CD44 between aFIB1 and MΦ1, may facilitate immune cell migration and promote differentiation toward a pro-inflammatory profile (*49, 50*). Additionally, the VEGFA-NRP2 LR pair between PT4 and MΦ2 may contribute to upregulating VEGFA signaling (*51*). Notably, VEGFA supplementation in the early phase of AKI has been shown to protect against acute kidney injury (*52*).

In post-KT AKI wo/Rec, the most prominent LR interactions were observed between MYOF and PT4. The FN1-SDC2 LR pair, identified between fibroblasts (aFB2, MYOF, pFIB) and PT4, may drive EMT (*53*). Additionally, strong interactions were detected between MYOF/mFIB/aFIB/aFIB21 and MΦ2, as well as between MYOF and MΦ1. The VCAM1-ITGA4_ITGB1 LR interaction between the injured PT4 cluster and immune cells (MΦ2/T) facilitates immune cell migration and differentiation into a pro-inflammatory phenotype (*50*). The SPP1-CD44 interaction between PT4 and immune cells (MΦ1/MΦ2/T/Treg) promotes neutrophil migration (*54*), leading to neutrophil accumulation in the interstitium, which increases vascular permeability, damages tubular and endothelial cells, and exacerbates kidney injury (*55*). The IL16-CD4 interaction between aFIB1 and Tregs suggests an IL16 role as a chemoattractant, facilitating Treg expansion (*56*). Notably, VEGFA-NRP2 interactions between aFIB1/aFIB2 and MΦ2, may represent a late-stage process, as aFIB1 cells likely originate from injured PT or StrProg in response to injury. Increased VEGFA levels in the late stage of AKI have been associated with fibrosis development (*52*).

### Co-culture assay validated the role of ischemic injury in promoting macrophage activation and fibroblast-like transitions

Hypoxic injury in PT cells induced marked transcriptional and phenotypic changes in monocyte-derived MΦs as evidenced in our co-culture assays (**Fig. 7A**). Hypoxic HK2 cells had significant upregulation of injury markers *HAVCR1, IL18*, and *VIM*. Concurrently, flow cytometry and qPCR analyses revealed elevated expression of α-SMA (fibroblast-associated marker), CD68 (MΦ-associated marker), TNFα (pro-inflammatory M1), and TGF-β (anti-inflammatory/pro-repair M2) in THP-1 monocytes co-cultured with hypoxic, injured HK2 cells compared to normoxic (non-injured) controls (**Fig. 7C–D**). These findings support the hypothesis that ischemic tubular injury drives MΦ activation toward a mixed M1/M2 polarization, potentially sustaining inflammation and fibrosis.

**Fig 7.**
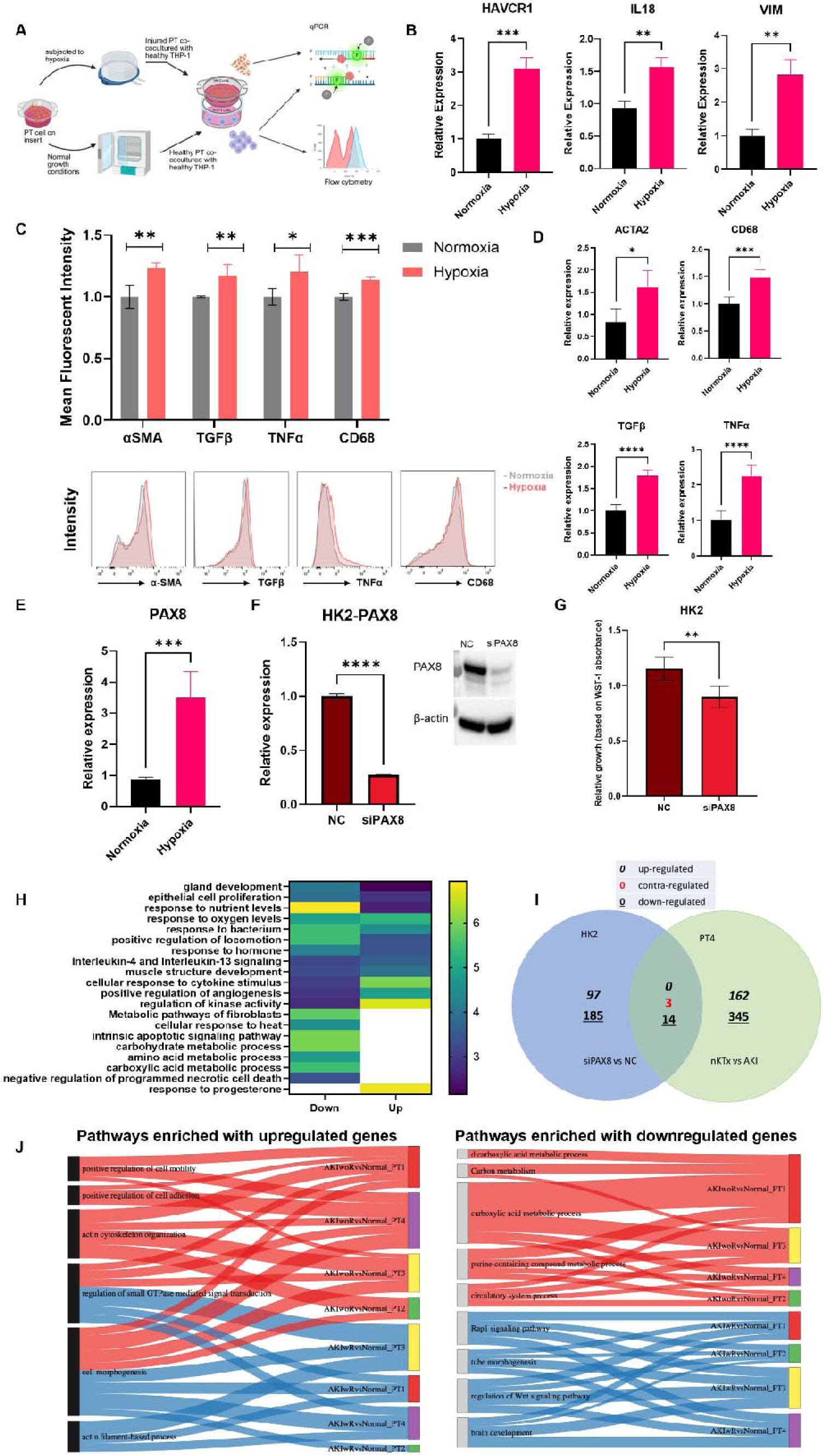
In vitro validations. (**A**) Co-culture methodology. (**B**) qPCR to quantify the upregulation of injury markers in injured HK2 cells. (**C**) Bar graph to show the mean fluorescent intensity for the indicated genes in THP-1 cells co-cultured with normoxic and hypoxic HK2 cells., bottom is the representative histogram. (**D**) qPCR for ACTA2, CD68, TGFβ and TNFα genes in co-cultured THP-1 cells (**E**) RNA-quantification by qPCR for PAX8 expression in injured HK2 cells. (**F**) qPCR (left) and western blot (right) image validating the knock down of PAX8. (**G**) WST-1 cell proliferation assay depicting decreased proliferation of HK2-siPAX8 cells. (**H**) Pathways enriched for the upregulated and downregulated genes in siPAX8 compared to HK2-NC. (**I**) Venn-diagram for the overlapping DEGs between HK2-siPAx8 v NC and PT4 DEGS nKTx vs AKI (Italics: upregulated genes; Underlined: downregulated; In red: contra-regulated). *p-*values for the bar graphs are *<0.05, **<0.005, ***<0.0005, ****<0.0001. (**J**) Sankey diagrams to visualize how bulk tissue pathways altered in DGF connect to cell type-specific changes in postKT-AKIw/Rec and postKT-AKIwo/Rec.

### *In vitro* PAX8 promotes epithelial proliferation and modulates injury-repair signaling

*PAX8* was expressed across all PT clusters (PT1, FC∼1.37; PT2, FC∼1.6; PT3, FC∼1.47; PT4∼FC1.89), with the highest upregulation observed in PT4 in the context of postKT-AKI, Critically, *PAX8* was significantly increased in injured HK2 cells (**Fig. 7E**), that were co-cultured with THP-1 monocytes mimicking findings from *ex vivo* human biopsies (**Fig. 3E**) suggesting its involvement in the injury response. To investigate its function *in vitro*, siRNA-mediated knockdown (KD) of PAX8 was performed in HK2 cells (siPAX8-HK2), with scrambled siRNA-transfected cells as controls (NC-HK2) (**Fig. 8F**). WST-1 assays revealed a ∼20% reduction in proliferation in siPAX8-HK2 cells at 48h, confirming a pro-proliferative role for PAX8 (**Fig. 8G**).

Bulk RNA-seq of siPAX8-HK2 cells identified DEGs involved in proliferation, injury, and repair (**Table S12**). Pathway analysis of downregulated genes showed enrichment in epithelial proliferation, apoptotic regulation, and protection against necrotic cell death, aligning with impaired growth upon PAX8 loss (**Fig. 8H**). Conversely, upregulated genes were enriched in cytokine response pathways, including IL-4 and IL-13 signaling, which promote M2 macrophage polarization and tissue repair.

Comparative analysis between DEGs from siPAX8-HK2 vs. NC-HK2 and PT4-nKT vs. AKI revealed 14 overlapping downregulated genes, including *HAVCR1, SPP1, CDH6*, and *FGFR1,* enriched in pathways related to EMT, angiogenesis, and KRAS signaling (**Fig. 8I, Table S13**). The shared downregulation of *HAVCR1*, a known injury marker, further supports the relevance of PAX8 in mediating tubular injury responses.

Together, these data position PAX8 as a key regulator of epithelial regeneration and immune signaling following AKI. Loss of PAX8 may impair adaptive repair and contribute to the pro-inflammatory, pro-fibrotic state observed in PT4 cells from postKT-AKIwo/Rec kidneys.

### Cross-platform validation of injury and repair signatures in DGF *versus* non-DGF kidney transplants and in DGF grafts with and without function recovery

DGF in KTRs is predominantly driven by IRI, which triggers epithelial damage and disrupts reparative processes. To validate the transcriptional programs identified by snRNA-seq, paired protocol biopsies (n = 90) performed at pre-implantation and 12 weeks post-transplant from DD KTRs were analyzed using microarrays, stratified by DGF (DGF vs. non-DGF; **Table 3**). Donor and recipient characteristics were well-matched, and no significant DEGs were detected at pre-implantation (FDR ≥ 0.102), indicating that post-transplant transcriptional changes reflect injury and repair processes rather than baseline differences. These findings further support the robustness of our snRNA-seq study design, which intentionally minimized baseline variability to focus on post-transplant injury and repair dynamics.

At12 weeks, 233 DEGs (FDR < 0.05, **Table S14**) distinguished DGF from non-DGF grafts. Pro-regenerative genes such as *EGF, FGF9,* and *FGFR2* were downregulated in DGF, alongside solute transporters (*KCNJ1, SLC12A1, SLC16A7*) and prostaglandin signaling (*PTGER3*), indicating epithelial dysfunction. The decline in expression of *ANGPT1* and *CD46* also reflected a compromised reparative microenvironment.

In contrast, DGF grafts exhibited upregulation of stress and fibrotic markers, including *DDX56, NGLY1, TNFRSF14, COL18A1,* and *LAMA5*, consistent with inflammation, proteotoxic stress, and maladaptive remodeling. Enhanced expression of autophagy-related *(ATG2A, ATG9A*) and mitochondrial dysfunction genes (*ME2, SUCLA2*) indicated cellular attempts at compensation. The up- and down-regulated genes identified in the microarray analysis were enriched in pathways that overlapped with those enriched in the PT snRNA-seq analysis. (**Fig. 7J**).

Sub-stratification of DGF grafts by long-term outcome (w/Rec vs. wo/Rec at 24 months) revealed enhanced suppression of regenerative and immune-regulatory pathways in non-recovering grafts (**Table S15**). These included *OSMR, SMAD5,* and *BMP2K* (tissue repair), *PTPRC* and *IL7R* (immune signaling), and proteostasis genes such as *HSPH1, TOP1,* and *FXR1*. Mitochondrial and cytoskeletal regulators (*NDUFA13, CALD1, GOLIM4*) were also reduced, suggesting structural disorganization and metabolic stress. Together, these results support a maladaptive transcriptional trajectory in DGF, especially in non-recovery cases, and validate the clinical and biological relevance of the epithelial, immune, and stromal injury states defined by snRNA-seq.

## Discussion

Ischemic AKI is the most frequent complication after kidney transplantation, significantly impairing both early and long-term graft function. This risk is further heightened in marginal kidneys, such as those from DCD donors, which are particularly susceptible to IRI. Despite its clinical relevance, the molecular mechanisms governing resolution versus progression to chronic dysfunction remain incompletely understood. This study addresses these gaps through the integration of snRNA-seq, IMC, and *in vitro* assays, revealing distinct epithelial, stromal, and immune trajectories that delineate recovery from maladaptive repair in human kidney allografts.

We leveraged a clinically relevant study design that captured biopsies within eight weeks post-transplant, identifying capturing key transitional states of injury response. Importantly, donor characteristics, including kidney quality, were comparable between recovery and non-recovery groups, minimizing confounding by baseline graft health. By integrating histological data and longitudinal graft function, we contextualized transcriptomic and spatial findings within clinically meaningful outcomes. The inclusion of both nKTx and NNK controls further enabled us to distinguish transplant-specific responses from baseline tissue states.

A central insight of this study is the critical role of (PT) cell fate in determining divergent outcomes in kidney grafts. We identified four distinct PT subclusters, representing a continuum from homeostasis (PT1), early adaptive response (PT2), transitional remodeling (PT3), to maladaptive dedifferentiation (PT4). Recovering grafts were enriched in PT3 cells, which exhibited transcriptional profiles associated with RNA metabolism, translation, and cellular homeostasis, features characteristic of adaptive repair pathways (*57, 58*). In contrast, PT4 cells, which predominate in non-recovering grafts, were marked by loss of epithelial identity, dedifferentiation, and activation of mesenchymal, pro-fibrotic, and immune-related gene programs. This phenotypic divergence aligns with the concept that epithelial plasticity, modulated by injury severity, local microenvironmental signals, and immune interactions, plays a decisive role in guiding whether the graft undergoes effective regeneration or progresses toward fibrosis (*59–63*).

A key novel finding of this study is the identification of regenerative STCs, characterized by co-expression of *PROM1*, *CD24*, and injury-associated markers, with significant enrichment in recovering grafts. While STCs have been described in animal models of kidney injury, their detection and expansion in human kidney allografts following AKI provide compelling evidence for their role in post-transplant epithelial repair (*64*). In humans, STCs are a heterogeneous and transient population that expands in response to injury, increasing in numbers with age and exposure to damaging stimuli (*64*). Prior studies have identified *CD24* and *CD133* (*PROM1*) progenitor-like cells in human kidneys capable of differentiating into tubular cells and podocytes (*65*). Our findings extend this knowledge by demonstrating that STC abundance is associated with graft recovery, suggesting that these cells actively contribute to epithelial regeneration. Therapeutic strategies aimed at enhancing STC expansion or function, such as administration of specific growth factors or small molecules, may offer a promising approach to improve post-transplant outcomes.

Conversely, non-recovering grafts were enriched in a unique population of tPT4 that exhibited a marked loss of epithelial identity and upregulation of fibroblast-associated genes, including multiple collagen isoforms (*COL1A1, COL5A1/2, COL6A1-3*) and mesenchymal regulators. These tPT4 cells represent a maladaptive epithelial state, poised between incomplete repair and fibrotic reprogramming. Transcriptional overlap between tPT4 and profibrotic fibroblast clusters (aFIB2, MYOF) suggests a continuum of epithelial-stromal transition, contributing to fibrogenesis. The emergence of tPT4 cells emphasizes the failure to restore epithelial integrity as a critical determinant of maladaptive repair and fibrosis, consistent with previous observations linking epithelial plasticity to graft outcomes (*66–68*).

Our pseudotime trajectory and L-R analyses reveal that epithelial, immune, and stromal cells are transcriptionally and spatially interconnected during graft injury and repair. In non-recovering grafts, PT4 cells exhibited strong interactions with profibrotic fibroblast subsets (aFIB2, MYOF) and pro-inflammatory macrophages (MΦ1), facilitating maladaptive remodeling. Key ligand-receptor pairs (VCAM1–ITGA4/ITGB2 and COL18A1–ITGA5) suggest that injured epithelial cells actively drive stromal and immune activation, beyond their role as passive injury targets. These interactions are consistent with known profibrotic signaling mechanisms and underscore epithelial plasticity as a determinant of fibrogenesis (*46, 52, 59–61, 69*).

We identified PAX8 as a transcription factor significantly upregulated during AKI, particularly within PT4 cells. Functional validation via PAX8 KD *in vitro* revealed impaired epithelial proliferation and increased pro-inflammatory signaling, suggesting that PAX8 promotes adaptive repair and modulates immune responses (*70–74*). These findings position PAX8 as a potential regulator of epithelial plasticity during injury. However, to confirm its therapeutic relevance, which is outside the scope of the current study, further assays, such as PAX8 overexpression, *in vivo* models, and direct epithelial-fibroblast co-culture systems, are warranted.

To address limitations associated with the relatively small snRNA-seq cohort, we validated our key findings using an independent, larger microarray dataset comprising deceased donor kidney transplant recipients stratified by DGF versus non-DGF outcomes. This cross-platform analysis confirmed central transcriptional programs identified by snRNA-seq, including the downregulation of regenerative and reparative genes (*EGF, FGF9, FGFR2, ANGPT1*) and solute transporters (*KCNJ1, SLC12A1, SLC16A7*), as well as upregulation of stress and fibrotic markers (*COL18A1, LAMA5, TNFRSF14, DDX56, NGLY1*), supporting the generalizability of epithelial dysfunction and fibrotic remodeling trajectories across molecular platforms.

Although previous studies in kidney transplantation have characterized AKI-related transcriptional changes using bulk microarray data (*19, 20*), our study extends these findings by delineating cell type–specific injury responses using high-resolution single-cell technology. Single-cell analyses in native kidney injury models (*75, 76*) have elucidated epithelial plasticity and maladaptive repair but lacked the complexity of allograft-specific ischemic and immune contexts. Our integrative approach—combining snRNA-seq, IMC, and functional assays (PAX8 KD, macrophage co-culture) using a unique set of well-characterized human graft samples, provides a robust framework for understanding maladaptive repair in the transplant setting. These findings establish a foundation for future multi-center studies to validate and expand upon these mechanistic insights.

This study presents the first high-resolution map of epithelial, immune, and stromal dynamics in human kidney allografts following AKI. We define transcriptional and spatial checkpoints that distinguish recovery from fibrosis and identify epithelial plasticity as a central determinant of graft fate. Therapeutic strategies aimed at preserving epithelial identity, modulating macrophage polarization, and disrupting pro-fibrotic signaling may offer promise for improving long-term transplant outcomes. Future multi-center studies and functional validations will be critical for translating these insights into clinical applications.

## Materials and Methods

### Patients and samples

This multicenter study included fourteen kidney samples from unique patients. Participants were enrolled in IRB-approved studies at two institutions: clinically indicated biopsies were collected at Montefiore, and surveillance biopsies at the University of Maryland Baltimore (UMB). Additional details are provided in the Supplementary Information. The study protocols were approved by the Institutional Review Boards of Montefiore/Einstein (IRB #09-06-174) and UMB (IRB #HP-00091954). All clinical and research activities were conducted in accordance with the ethical principles outlined in the Declaration of Istanbul on Organ Trafficking and Transplant Tourism.

### Sample processing and single-nucleus isolation

Nuclei isolation from tissue samples was performed using an adapted protocol with Nuclei EZ Lysis Buffer (NUC-101; Sigma-Aldrich), supplemented with protease (5892791001) and RNase inhibitors (N2615, Promega; AM2696, Life Technologies), as detailed in the Supplementary Information.

### Library preparation and quality control

Library preparation began with single-nucleus lysis, followed by reverse transcription of RNA within each droplet into complementary DNA (cDNA) using the 10x Chromium Single Cell 5′ Library & Gel Bead Kit v2 (10x Genomics), according to the manufacturer’s instructions and as detailed in the Supplementary Information.

### snRNA-seq data analysis

FastQ files generated by the 10x Genomics sequencing pipeline were aligned to the human pre-mRNA reference genome (GRCh38) using CellRanger (v3, 10x Genomics) and analyzed with the Seurat R package, as described in the Supplementary Information.

### Gene ontology analysis

Gene set enrichment analysis was done as described in the Supplementary file.

### Ligand receptor analysis

Cell–cell interaction analysis was performed separately for each of the three conditions using LIANA, as described in the Supplementary Information.

### Single cell trajectory analysis

Single-cell pseudotime trajectory dynamics were performed using Monocle version 3.0, as described in the Supplementary Information.

### Cell origin determination

To determine the origin of cells in the allograft, we leveraged sex mismatches between kidney transplant recipients (KTRs) and their donors. Donor-versus recipient-derived cells were identified based on the expression of the Y chromosome-encoded *KDM5D* and the X chromosome-encoded *XIST*. Cell clusters were first separated by individual sample, and origin was inferred based on sex-specific gene expression profiles.

### Imaging Mass Cytometry

Imaging mass cytometry (IMC) was performed at the Histopathology and Tissue Shared Resources Core of Georgetown University using formalin fixed, paraffin-embedded tissue sections. The antibody panel is provided in the Supplementary Information.

### Cell culture

Human kidney epithelial cells (HK2, ATCC Cat no: CRL-2190) and human monocyte cells (THP-1, ATCC Cat no: TIB-202) were cultured according to the manufacturer’s instructions.

### Co-culture assay

For co-culture experiments were performed using HK-2 and THP-1 (ATCC, Cat no: TIB-202) were used, as described in the Supplementary file.

### Flow cytometric analysis

THP-1 cells were analyzed by flow cytometry, following the protocol described in the Supplementary Information.

### siRNA knockdown of transcription factor candidates

Knockdown of transcription factor genes in renal proximal tubular epithelial cells (RPTECs) was performed using Invitrogen’s siRNA transfection protocol, as described in the Supplementary Information.

### WST-1 Assay for cell proliferation and viability

The WST-1 assay was conducted using Roche’s protocol, as described in Supplementary Information.

### qPCR

Total RNA was isolated from THP-1 cells using the Quick-RNA Mini Prep Kit (Zymo Research, Cat. No. R1055), and reverse transcription was performed with the iScript Reverse Transcription Supermix (Bio-Rad, Cat. No. 1708840), according to the manufacturer’s instructions. Gene expression was quantified using TaqMan probes targeting TGFβ, ACTA2 (α-SMA), TNF-α and CD68.

### Data Statement

The data supporting the findings of this study will be made publicly available (GEO ID: GSE284634 and GSE269078). Affymetrix GeneChip microarrays (HG-U133A 2.0) are accessible publicly using GEO ID: GSE147451

## Abbreviations

PostKT-non-DGF: post kidney transplantation without delayed graft function;
PostKT-DGF: post kidney transplantation with delayed graft function; post kidney transplantation without delayed graft function
w/Rec: with functional recovery
wo/Rev: without functional recovery
DCD: donation after circulatory death
ESRD: end stage renal disease
KDPI: Kidney Donor Profile Index
DM/HTN: Diabetes Mellitus/Hypertension
DM: Diabetes Mellitus
HTN: Hypertension
FSGS: Focal segmental glomerulosclerosis
A: Asian
AA: African American
C: Caucasian
H: Hispanic
O: Other
eGFR: estimated glomerular filtration rate measured in mL/min/1.73m².

## Acknowledgements

This research reported in this publication was supported by the National Institutes of Health R21DK100678 (DGM); National Institutes of Health R01DK109581 (VRM); National Institutes of Health R01DK122682 (VRM); National Institutes of Health R21AI172077 (VRM). We would also like to thank the members of the Institute for Genome Sciences at the University of Maryland, Baltimore, for their invaluable contribution and assistance using the 10x Genomics Chromium.

## Author contributions

Conceptualization: VRM, EA, DGM, SA, TVR, HZ

Methodology: TVR, SA, HZ, ACS

Investigation: VRM, SA, TVR, HZ, ACS, EA, DGM

Clinical data acquisition: VRM, DGM, EA, SM

Visualization: SA, TVR, HZ, ACS, CD

Funding acquisition: VRM, DGM

Project administration: VRM, DGM, EA

Supervision: VRM, EA, DGM

Writing – original draft: VRM, SA, TVR, HZ

Writing – review & editing: VRM, SA, TVR, HZ, ACS, EA, DGM, CML, AR, SM, JB, MM

## Supplemental Information Supplementary Tables

**Table S1.** Summary of Illumina sequencing results

**Table S2.** Summary of Q30 scores.

**Table S3.** Counts of isolated cells in each cell cluster.

**Table S4.** Gene-based markers for cell clusters identification.

**Table S5.** List of genes used for the classification of cluster identity

**Table S6.** List of differentially expressed genes between NNK versus nKTx

**Table S7.** Differentially expressed genes between AKI versus nKTx

**Table S8.** Fibroblast cell type markers

**Table S9.** Immune cell type markers.filtered.txt

**Table S10.** Differentially expressed genes between postKTw/Rec versus postKTwo/Rec

**Table S11.** Ligand receptor interactions

**Table S12.** Differential expression analysis table HK2 siPAX8

**Table S13.** Pathways enriched for 14mutual downregulated genes in siPAX8-HK2 vs NC-HK2 and PT4-nKT vs AKI

**Table S14.** Differentially expressed genes comparing postKT-wDGF versus postKT-wo DGF cross-sectionally at 12 weeks

**Table S15.** Differentially expressed genes between postKT-wDGF w/Rec and postKT-woDGF wo/Rec

## Supplementary Figures

**Fig S1.** Quality control (QC) parameters assessed

**Fig S2.** (A) UMAP demonstrating all identified clusters and the integration of samples and clusters between the four conditions. (B) Cell portion in nKTX, NNK, postKT-AKIw/Rec and postKT-AKIwo/Rec

**Fig S3.** Enriched pathways associated with the integrated genes for TC cluster

**Fig S4.** UMAP and pseudotime trajectory analysis on integrated proximal tubule (PT) and fibroblast (Fib) subclusters across conditions

**Fig S5.** Ligand-receptor interaction in native normal kidneys (NNK) and normal kidney transplants (nKTx)

## Notes

### Competing Interest Statement

The authors have declared no competing interest.

