## Supplemental Information for "Epithelial-Immune-Stromal Interactions Define Divergent Repair and Fibrosis Pathways After Acute Kidney Injury in Human Renal Transplants"

**Valeria R. Mas, MSc, PhD, FAST**

Division of Surgical Sciences, Department of Surgery, University of Maryland School of Medicine, 670 W Baltimore Street, HSFIII Building, Room 7179, Baltimore, Maryland 21201, USA.

**List of Supplemental Information**

**Supplementary Tables (present in ‘Supplementary Information.xlsx’ file)**

Table S1. Summary of Illumina sequencing results

Table S2. Summary of Q30 scores.

Table S3. Counts of isolated cells in each cell cluster.

Table S4. Gene-based markers for cell clusters identification.

Table S5. List of genes used for the classification of cluster identity

Table S6. List of differentially expressed genes between NNK versus nKTx

Table S7. Differentially expressed genes between AKI versus nKTx

Table S8. Fibroblast cell type markers

Table S9. Immune cell type markers.filtered.txt

Table S10. Differentially expressed genes between postKTw/Rec versus postKTwo/Rec

Table S11. Ligand receptor interactions

Table S12. Differential expression analysis table HK2 siPAX8

Table S13. Pathways enriched for 14mutual downregulated genes in siPAX8-HK2 vs NC-HK2 and PT4-nKT vs AKI

Table S14. Differentially expressed genes comparing postKT-wDGF versus postKT-wo DGF cross-sectionally at 12 weeks

Table S15. Differentially expressed genes between postKT-wDGF w/Rec and postKT-woDGF wo/Rec

**Supplementary Figures**

Fig S1. Quality control (QC) parameters assessed

Fig S2. (A) UMAP demonstrating all identified clusters and the integration of samples and clusters between the four conditions. (B) Cell portion in nKTX, NNK, postKT-AKIw/Rec and postKT-AKIwo/Rec

Fig S3. Enriched pathways associated with the integrated genes for TC cluster

Fig S4. UMAP and pseudotime trajectory analysis on integrated proximal tubule (PT) and fibroblast (Fib) subclusters across conditions

Fig S5. Ligand-receptor interaction in native normal kidneys (NNK) and normal kidney transplants (nKTx)

**Supplementary Materials and Methods**

**Patients and samples**

Kidney biopsies from eleven deceased donor kidney transplant recipients (KTRs) and three biopsies from living donors (normal native kidneys, NNK) were studied. From 14 KTR graft biopsies, four kidney grafts with normal/stable graft function (nKTx), whose biopsies were categorized as normal/non-specific, and seven AKI posttransplant biopsies (postKT-AKI) (collected within 8 weeks post-KT). nKTx were >15 months post-transplantation, had an estimated GFR (eGFR) of ≥60 mL/min/ 1.73m^2^, no proteinuria, no circulating IgG antibodies against donor HLA at the time of biopsy, and had normal/non-specific findings in the allograft surveillance biopsies. Kidney allograft tissue was obtained using an 18-gauge biopsy needle and all samples were immersed in RNAlater (Ambion) immediately after collection. Clinical data was collected on all patients by chart review. All patients received triple-drug immunosuppression that included tacrolimus, mycophenolate mofetil, and prednisone. Biopsies were examined by light microscopy using hematoxylin and eosin, periodic acid-Schiff (PAS), Masson Trichrome and C4d immunoperoxidase stains. Evaluation of the biopsies was based on the Banff acute and chronic indices including glomerulitis (g), interstitial inflammation (i), tubulitis (t), intimal arteritis (v), peritubular capillaritis (ptc), transplant glomerulopathy (cg), mesangial matrix increase (mm), interstitial fibrosis (ci), tubular atrophy (ct), and vascular fibrous intimal thickening *(1, 2)*. Estimated glomerular filtration rate (eGFR) was calculated using the 2021 CKD-EPI equation, which incorporates serum creatinine, patient age, and sex to provide an accurate assessment of renal function. This updated equation eliminates race-based coefficients, aligning with current best practices for equitable clinical assessment. The eGFR values derived were used to evaluate kidney function in the study population and to correlate renal outcomes with clinical and experimental variables.

A graft was considered to have recovered function if it demonstrated a continuous positive slope in eGFR following transplantation, with an eGFR of ≥50 mL/min/1.73 m² at 24 months post-transplant. This definition reflects sustained improvement in renal function over time and meets clinically meaningful thresholds for graft performance. Grafts classified as non-recovery were characterized by a failure to sustain a positive trajectory in eGFR post-transplant, with values remaining below 50 mL/min/1.73 m² at 24 months *(3, 4)*. In addition, for-cause biopsies performed after the first 4 weeks post-transplant revealed increased interstitial fibrosis and tubular atrophy (IFTA), along with other chronic lesions as defined by the Banff Classification. IFTA, indicative of chronic allograft injury, was graded based on the percentage of cortical involvement, with Grade I *(1, 5, 6)* involving 6–25%, Grade II 26–50%, and Grade III greater than 50% of the cortical area. Chronic lesions also included features such as chronic glomerulopathy (cg), arterial intimal fibrosis (cv), and transplant glomerulopathy, reflecting ongoing immune and non-immune damage.

Delayed graft function (DGF) was defined as ischemic acute kidney injury (AKI) based on the requirement *(7–10)* for dialysis within the first 7 days post-transplant, without evidence of rejection or other non-ischemic causes, in cases where biopsies were not performed. Inclusion criteria for DGF classification required: (i) initiation of dialysis within 7 days post-transplant, (ii) absence of clinical or laboratory signs suggestive of hyperacute or acute rejection, (iii) no biopsy-confirmed alternative diagnosis, and (iv) a clinical context consistent with ischemia-reperfusion injury, including prolonged cold ischemia time or donor factors associated with increased ischemic risk. This definition ensured that DGF was attributed specifically to ischemic AKI in the absence of confounding pathological findings.

A total of 45 kidney transplant recipients were included in the analysis, all of whom had protocol biopsies performed at pre-implantation and again at 12 weeks post-transplant. Patients were stratified based on the presence or absence of delayed graft function (DGF), with 29 in the postKTx-non-DGF group and 16 in the postKTx-DGF group. Recovery of graft function occurred in 69.0% of the non-DGF group and 62.5% of the DGF group (p = 0.746).

There were no statistically significant differences between groups in donor age, sex, cause of death, kidney donor profile index (KDPI), cold ischemia time (CIT), or other major donor characteristics. However, recipients in the DGF group had significantly higher body weight and BMI (95.67 vs. 75.90 kg, p < 0.001; 30.92 vs. 26.38 kg/m², p = 0.004), and greater recipient height (176.38 vs. 169.14 cm, p = 0.014). Early post-transplant renal function was significantly lower in the DGF group, with reduced eGFR at 1 week (12.48 vs. 51.15 mL/min/1.73 m², p < 0.001), 1 month (38.90 vs. 64.22, p < 0.001), and at multiple later time points including 3-, 6-, 15-, and 24-months post-transplant (all p < 0.05). The DGF group also had higher serum creatinine levels on days 1 and 2 post-transplant (p = 0.031 and 0.026, respectively) and lower creatinine reduction ratios, although this did not reach statistical significance.

Of the 16 patients with DGF, only 7 underwent for-cause biopsies within the first 6 weeks post-transplant, all of which demonstrated acute tubular necrosis (ATN) without evidence of additional acute or chronic lesions. Among the 16 DGF patients, 12 showed clinical recovery of graft function based on eGFR trajectories; however, at 24 months post-transplant, all of them had eGFR values below 50 mL/min/1.73 m², ranging from 28 to 50 mL/min/1.73 m², indicating persistent suboptimal long-term graft function despite initial recovery.

Overall, this cohort allows longitudinal assessment of graft function and histologic progression in the context of DGF, providing a foundation for evaluating molecular and cellular pathways associated with graft recovery versus non-recovery.

**Sample processing and single nuclei isolation**

Nuclei isolation from tissue samples followed an adapted protocol utilizing Nuclei EZ Lysis buffer (NUC-101; Sigma-Aldrich), supplemented with protease (5892791001) and RNase inhibitors (N2615, Promega; AM2696, Life Technologies). Briefly, tissue samples were mixed with 2 mL of ice-cold lysis buffer, incubated on ice for 5 minutes, and then combined with an additional 2 mL of buffer. The lysate was then filtered through a 40 µm cell strainer (43-50040-51; pluriSelect) before centrifugation at 500 x g for 5 minutes at 4°C. After discarding the supernatant, the pellet was resuspended in fresh buffer, incubated on ice for another 5 minutes, and centrifuged again. The final pellet was resuspended in Nuclei Suspension Buffer (PN-2000153; 10x Genomics; comprising 1x PBS, 1% bovine serum albumin, and 0.1% RNase inhibitor), followed by additional filtration. DAPI staining enabled visualization, and nuclei counts were obtained using the Countess 3 Automated Cell Counter (ThermoFisher). Finally, the 10x Chromium system (10x Genomics) was used to encapsulate viable nuclei into droplets with barcode-linked gel beads.

**Library preparation and quality control**

Library preparation began with single nuclei lysis, where RNAs within each droplet were reverse transcribed into complementary DNA (cDNA) using the 10x Chromium Single Cell 5′ Library & Gel Bead Kit (v2) (10x Genomics), following the manufacturer’s guidelines. After emulsion breaking, cDNA underwent amplification, fragmentation, and adapter ligation, using the same kit as per 10x Genomics protocols. Thermal cycling conditions were set at 98°C for 45 seconds, followed by 16 cycles of 98°C for 20 seconds, 67°C for 30 seconds, and 72°C for 1 minute, ending with a 72°C hold for 1 minute and then held at 4°C. Post-PCR, cDNA concentration was measured with a Qubit fluorometer (ThermoFisher). A Bioanalyzer (Agilent) and gel electrophoresis were employed to confirm expected fragment sizes after adapter addition. Libraries were then multiplexed, pooled, and sequenced on a NovaSeq 6000 S4 flow cell (Illumina) using 150 bp paired-end sequencing.

**snRNA-seq data analysis**

FastQ files produced by the 10x Genomics sequencing pipeline were aligned to the human pre-mRNA reference genome (GRCh38) using CellRanger (v3, 10x Genomics). This alignment yielded three primary output files per sample: a list of cell barcodes, a gene name table, and a gene expression matrix. Initial quality control (QC) on CellRanger outputs involved evaluating feature counts, gene expression depth, and the percentage of mitochondrial gene expression in each kidney sample. Genes detected in fewer than three cells, cells with low gene counts (fewer than 400 genes), or cells with excessive gene expression (>5,000 genes) were filtered based on data-informed thresholds. Additionally, nuclei with over 2.5% mitochondrial gene expression were excluded from further analysis.

Samples were combined into a single dataset using the ‘Seurat’ R package, followed by checks for batch effects due to technical variations or cell cycle influences on proliferative states. Batch effect correction was followed by cell clustering, which was carried out via principal component analysis (PCA) and uniform manifold approximation and projection (UMAP), revealing clusters based on gene expression profiles. Expression patterns across clusters were then analyzed to identify unique marker genes for each cluster, which were cross-referenced with cell-type-specific genes from public databases through gene set enrichment analysis to infer likely cell identities.

We identified primary cell types across various conditions and found uncharacterized cells marked by non-specific markers. Cell type proportions were then compared between conditions to observe population differences. Further downstream analyses involved sub-clustering of relevant cell types, determining differentially expressed genes (DEGs) within each type and subtype under different graft conditions. DEGs with FDR ≤ 0.05 and fold change ≥ 1.5 were used for gene ontology and pathway enrichment analysis. All statistical analyses and visualizations were performed in R.

**Gene ontology analysis**

Gene set enrichment analysis for intra- and inter-cluster comparisons across study groups (normal and fibrosis) was conducted using Enrichr *(11)* and Metascape *(12)*, with a significance threshold of FDR ≤ 0.05 to identify enriched pathways. Differentially expressed gene (DEG) lists were also uploaded to WebGestalt *(13)* and the DAVID Bioinformatics Resource (v6.8) *(14)* for annotation, linking these genes to pertinent biological functions.

**Ligand receptor analysis**

To assess potential cell-cell interactions between all cell types separately for each of the three conditions, we utilized LIANA, LIgand-receptor ANalysis framework interface. LIANA integrates multiple LR-algorithms and compiles a consensus score utilizing a Robust Rank Aggregate method *(15)*. The consensus score is interpreted as adjusted p-values and, thus, LR pairs with consensus scores <0.01 were considered for downstream analysis. Visualizations were generated using R graphical software.

**Single cell trajectory analysis**

Single cell dynamics were observed using Monocle 3 *(16)*. Monocle is an unsupervised algorithm that temporally orders single-cell expression profiles in pseudotime and allows for multiple cell fates from a single progenitor cell type (as indicated by multiple branching events) *(16–18)*. For each condition, the Seurat SummarizedExperiment object was transformed into the required CellDataSet object utilized as input for Monocle3. The gene expression matrix was used to determine cell clusters that were further grouped into cell partitions. For a predetermined set of cell types, trajectory graphs were inferred using the gene expression changes to place cells along a pseudo timeline as they transition from one state to the next. Based on the learned trajectories, monocle was further utilized to model changes in gene expression as a function of pseudotime. Significant pseudotime-related genes were selected using an adjusted p-value < 5%, Moran’s I statistic > 0, and genes expressed in at least 10% cells. Visualizations were generated using R graphical software.

**Imaging mass cytometry**

Marker panels were ordered from Fluidigm (Standard Biotool) which included α-smooth muscle actin, (αSMA, catalog #3141017D), CD4 (catalog #3156033D), CD68 (catalog #3159035D), Collagen 1A (COL1A1, catalog #3169023D), E-cadherin (ECAD, catalog #3158029D), and Vimentin (VIM, catalog #3143027D). Aquaporin 1 (AQP1) was purchased from AbCam (catalog #AB168387). Images were obtained using the Fluidigm Hyperion Imaging System with appropriate fluorescence filters and processed using the Quantitative Pathology and Bioimage Analysis (QuPath) software v0.3.2.S15. Control experiments were performed and yielded no observable non-nonspecific staining. All patient samples were deidentified prior to imaging.

Quantification of fibrotic markers (COL1A1, αSMA, and VIM) from imaging mass cytometry (IMC) data was performed using ImageJ (NIH, USA). For each marker, images were first split into individual grayscale channels corresponding to COL1A1 (white), αSMA (green), and VIM (yellow). Consistent threshold values were applied across all images to segment positive staining from background, using the “Adjust Threshold” function. Binary masks were generated to identify areas of positive signal, and the percentage of area positive for each marker was calculated using the “Analyze Particles” function, expressed as area fraction (% positive area per total field of view). Multiple ROIs were analyzed within a single tissue section, and the variation across those ROIs was used to compute mean ± SD for each condition. All quantifications were performed under blinded conditions to ensure unbiased measurement. Also, biological replication for IMC quantification was performed for postKT-AKI wo/Rec (n=3 samples).

To assess baseline fibrotic remodeling in stable transplant grafts, we quantified the expression of COL1A1, vimentin (VIM), and α-smooth muscle actin (αSMA) in nKTx samples IMC. Minimal fibrotic marker expression was detected in nKTx grafts. Specifically, the mean area of COL1A1+ staining was 1.3% ± 0.5%, VIM+ area constituted 2.1% ± 0.6%, and αSMA+ area was limited to 1.0% ± 0.4% of the total tissue section. These results confirm a homeostatic epithelial and stromal environment in stable allografts, characterized by preserved tubular architecture (AQP1+, eCAD+) and low fibrotic marker expression.

We next evaluated fibrotic remodeling in PostKT-AKIw/Rec allografts. IMC analysis demonstrated a moderate increase in fibrotic markers compared to stable grafts. The mean area of COL1A1+ staining was 5.6% ± 1.2%, VIM+ area was 7.3% ± 1.5%, and αSMA+ area reached 4.5% ± 1.1% of total tissue area. Spatial distribution revealed focal expression of COL1A1 and αSMA predominantly in peritubular regions, with partial retention of epithelial architecture (AQP1+, eCAD+). These findings indicate localized fibrotic remodeling during recovery, in contrast to diffuse fibrosis observed in non-recovering grafts.

Kidney allografts that failed to recover function (PostKT-AKIwo/Rec) exhibited pronounced fibrotic remodeling, as demonstrated by IMC analysis. The mean COL1A1+ area was markedly elevated at 15.2% ± 2.1%, VIM+ area reached 18.7% ± 2.5%, and αSMA+ area was 11.4% ± 1.8% of total tissue area. These fibrotic markers displayed diffuse distribution, particularly in regions of disrupted epithelial integrity, evidenced by reduced AQP1 and eCAD staining. The spatial co-localization of COL1A1, VIM, and αSMA confirmed extensive mesenchymal activation and extracellular matrix deposition, consistent with maladaptive repair.

**Co-culture protocol**

For co-culture experiments, human kidney cells HK2 and the human monocyte cell line THP-1 (ATCC, Cat no: TIB-202) were used. HK2 cells (0.5 million) were seeded on 1µm pore size inserts (Falcon, Cat no: 353102), while 0.2 million THP-1 cells were plated in a 6-well plate. After 16 hours, the HK2 cells in the inserts were subjected to 45 minutes of 95% nitrogen gas balanced with 5% CO_2_ to mimic hypoxia and then transferred under hypoxic conditions to 4°C for 6 hours. Control HK2 cells in inserts were maintained under normoxic conditions (37°C and 5% CO_2_). Following hypoxic exposure, the inserts were transferred to the THP-1 plates and co- incubated for 72 hours at 37°C and 5% CO_2_. Subsequently, co-cultured THP-1 cells were isolated for flow cytometry analysis and RNA isolations for qPCR analysis.

**Flow cytometric analysis**

For flow cytometry, co-cultured-treated THP-1 cells were stimulated with ionomycin and phorbol 12-myristate 13-acetate, followed by addition of Brefeldin (GolgiPlug, BD Biosciences, Cat no: 555029), and Monensin (GolgiStop, BD Biosciences, Cat no: 554724) after 1 hour. Cells were then incubated for an additional 4 hour at 37°C to allow for intracellular cytokine accumulation and staining. They were then stained for viability using eFluor 780 Fixable Viability Dye (eBioscience, Cat no: 65-0865-14)) and blocked using Human Fc Block (BD Biosciences, Cat no: 564219) for 10 minutes at 4°C in Phosphate-buffered saline. Extracellular staining was done for CD163 APC (Biolegend, Cat no: 326510) and CD14 BV510 (Biolegend, Cat no: 123323) for 20 minutes at 4°C in FACS buffer. After extracellular staining, cells were washed with FACS buffer and treated with FOXP3/Transcription Factor staining buffer kit (eBioscience) for intracellular staining. Cells were intracellularly stained with CD68 PerCP-Cy5.5 (Biolegend, Cat no: 333814), TGF-β PE (BD Biosciences, Cat no: 562339), TNF-α PE-Cy7 (Invitrogen, Cat no: 25-7349-82), and α-SMA V450 (Novus Biologicals, NBP2-59429MFV450) for 30 minutes at 4°C. Following staining, cells were washed with FACS buffer and resuspended for data collection. Cells were collected on a BD LSRFortessa using BD FACSDiva software and analyzed in FlowJo (BD Biosciences). Data was transferred to GraphPad Prism 10 for graphical illustration and statistical analysis.

**siRNA knockdown of TF candidates in human renal proximal tubular epithelial cells**

Invitrogen siRNAs (Negative Control 1 (NC1): Cat no: AM4611, siPAX8 (Cat no AM16708), were diluted and used to transfect the cells (HK2) using Lipofectamine RNAiMAX (Thermo Fisher, 13778075) following the manufacturer’s protocol. RNA and protein were extracted after 48h. qPCR was done using a PAX8 TaqMan probe, (*PAX8*: Assay ID: Hs00247586m1, Cat no: 4331182). Relative expression was calculated with respect to the expression of housekeeping gene *RPL27* (Assay ID: Hs03044961g1, Cat no: 4331182). Student t-test was done to compare non-targeting and siRNA treatment groups (p < 0.05). Western blot was done to confirm the knock-down of *PAX8* at protein level using the Invitrogen monoclonal antibody (INV-MA1117). Bulk RNA-seq was done to evaluate expression of target genes and to determine the changes in transcriptome upon the knockdown of PAX8. Differentially expressed genes (p-value <0.05 and |log_2_FC| >0.5 were then subjected to further analysis using Enrichr *(11)*, Metascape *(12)* and ShinyGO *(19)* gene-set enrichment tools.

**WST-1 Assay protocol for cell proliferation and Viability**

HK2 cells (5000 cell count) were seeded in a 96 well-plate and treated with the siRNAs NC1 and siPAX8, as mentioned above. The cells were incubated for 48h at 37°C and 5% CO_2_. 90µl of fresh media was added to the wells. For the assay, 10µl of cell proliferation reagent WST-1 (Roche, Cat no: 05015944001) was added to the plate and incubated for 4h at 37°C and 5% CO_2_. After shaking the plate at 37°C for an hour, absorbance was measured at 480nm.

**Microarray data analysis**

The Affymetrix GeneChip HG-U133A 2.0 was read into the R programming environment using the affy *(20)* Bioconductor package. As a QC measure, the MAS 5.0 expression summaries were calculated to form the 3’:5’ ratios for GAPDH. Probe set expression summaries were obtained using the robust multiarray average method. The Affymetrix Detection Call algorithm was used to determine whether probe sets were present, marginally present, or absent in each sample. The gene expression data matrix was then filtered to exclude control probe sets and probe sets called absent in all samples, leaving 19,293 probe sets for statistical analysis.

We performed a cross-sectional analysis comparing post-KT samples at each time point with respect to postKTx-DGF (with *versus* without) using a moderated t-test with adjustment for multiple hypothesis tests using the false discovery rate.

Pathway analysis comparing kidney allograft tissue with and without Delayed Graft Function (DGF) revealed significant molecular signatures that correlate with distinct recovery patterns observed in our single-nucleus RNA-sequencing data. The Sankey diagrams visualize how bulk tissue pathways altered in DGF connect to cell type-specific changes in AKI with recovery (AKIwR) and AKI without recovery (AKIwoR) conditions.

Pathways downregulated in DGF allografts showed distinct patterns of association with recovery status. Metabolic pathways including dicarboxylic acid metabolism, carbon metabolism, carboxylic acid metabolism, and purine-containing compound metabolism predominantly connected to the AKIwoR condition, suggesting these metabolic alterations may be critical determinants of non-recovery outcomes. Circulatory system processes also showed strong connections to non-recovery states. In contrast, developmental and signaling pathways including Rap1 signaling, tube morphogenesis, Wnt signaling regulation, and brain development connected primarily to the AKIwR condition, indicating these pathways may facilitate recovery following kidney injury.

The upregulated pathway analysis revealed complementary insights. Pathways promoting cell motility, adhesion, actin cytoskeleton organization, and small GTPase-mediated signal transduction were predominantly associated with the AKIwoR condition. This suggests excessive cell adhesion and motility responses may contribute to maladaptive outcomes. Conversely, cell morphogenesis and actin filament-based processes connected primarily to the AKIwR condition, potentially representing adaptive repair mechanisms that facilitate recovery.

The analysis also demonstrated proximal tubule subtype-specific responses, with PT1 and PT4 showing strong connections to adhesion and motility pathways in non-recovery states, while PT3 exhibited prominent associations with morphogenic pathways in recovery states. These findings suggest that specific PT subtypes may drive distinct aspects of the injury response, with potential implications for targeted therapeutic approaches to promote recovery following AKI.

These results provide a molecular framework connecting bulk tissue alterations in DGF to cell type-specific recovery mechanisms, highlighting potential determinants of recovery versus non-recovery outcomes in acute kidney injury.

**Supplementary Figures**

**Fig S1:** Quality control (QC) parameters assessed: all the samples passed QC and 38,534 high-quality nuclei were retained for further analysis.

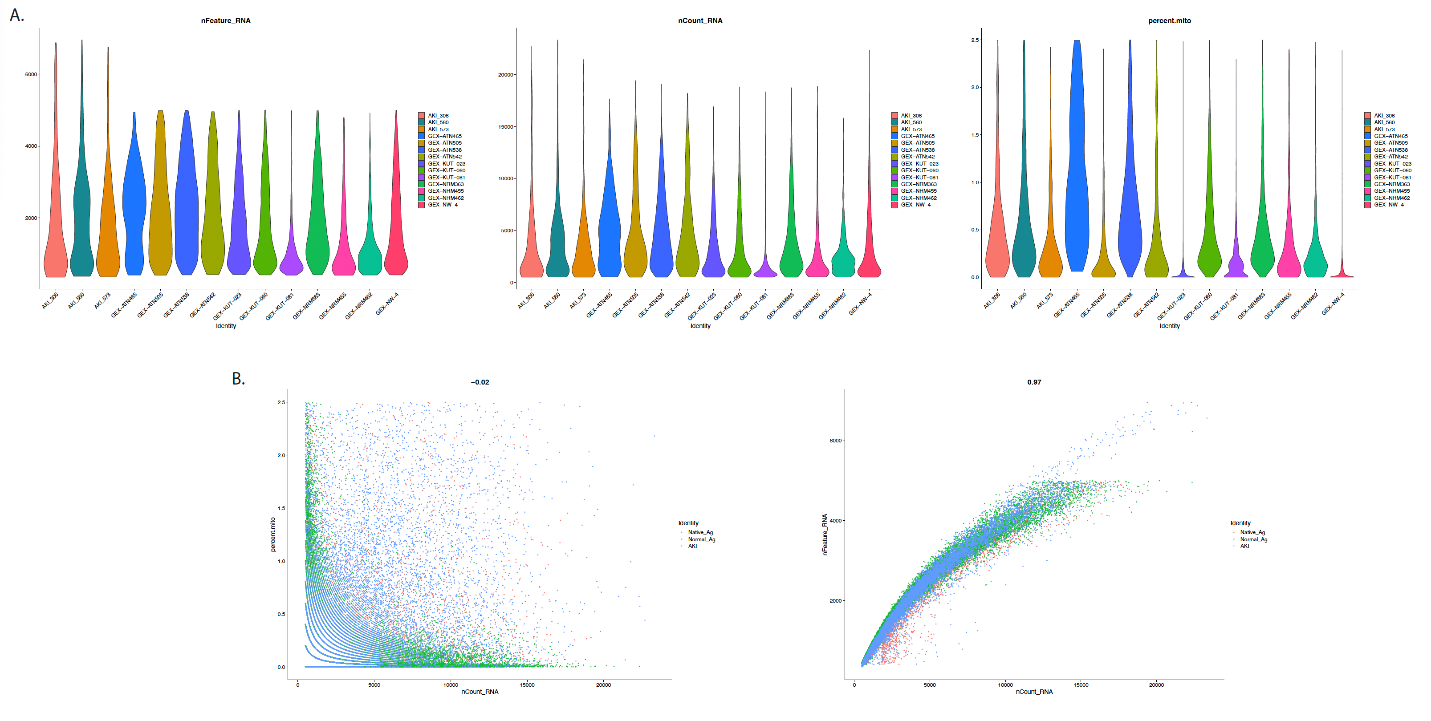

**Fig S2 (A)** UMAP demonstrating all identified clusters and the integration of samples and clusters between the four conditions. (B) Cell portion in nKTX, NNK, postKT-AKIw/Rec and postKT-AKIwo/Rec

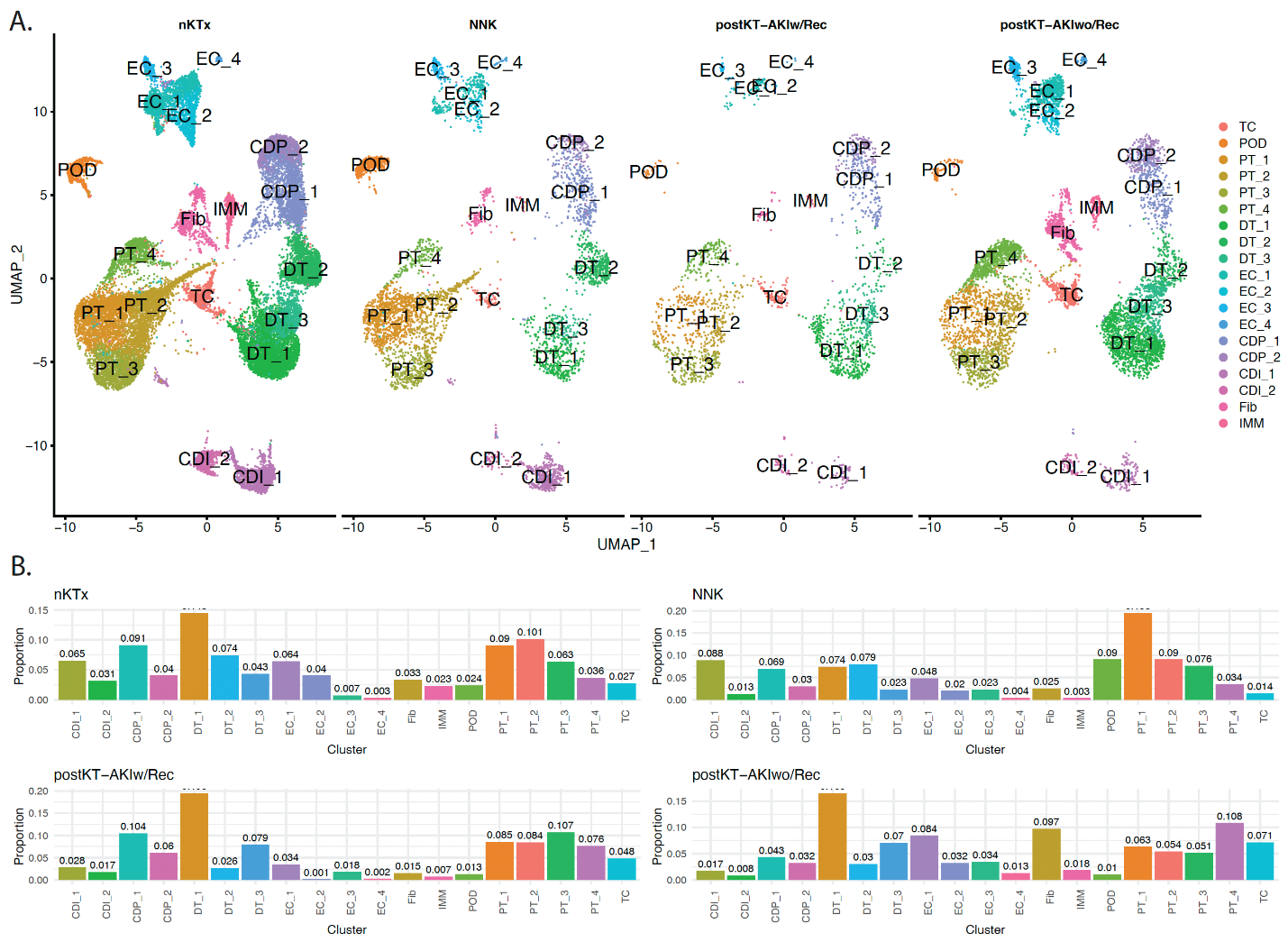

**Fig S3:** Enriched pathways associated with the integrated genes for TC cluster

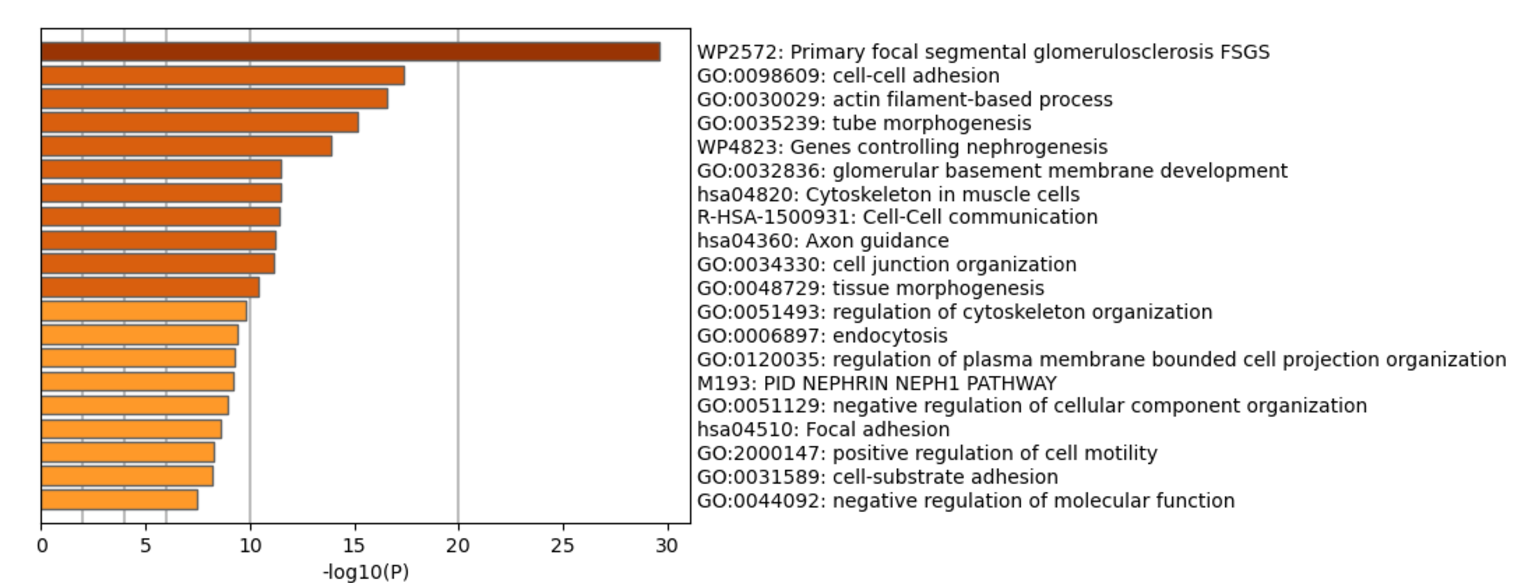

**Fig S4: Integrated clustering of PT and Fib populations across conditions**

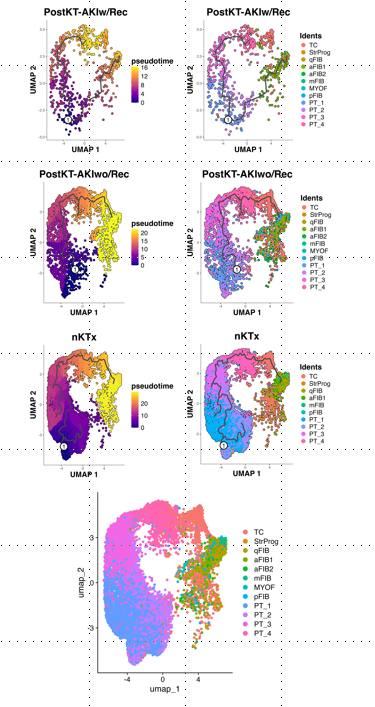

**Fig S5:** Ligand-receptor interactions between PT and Fibroblasts in NNK and nKTx.

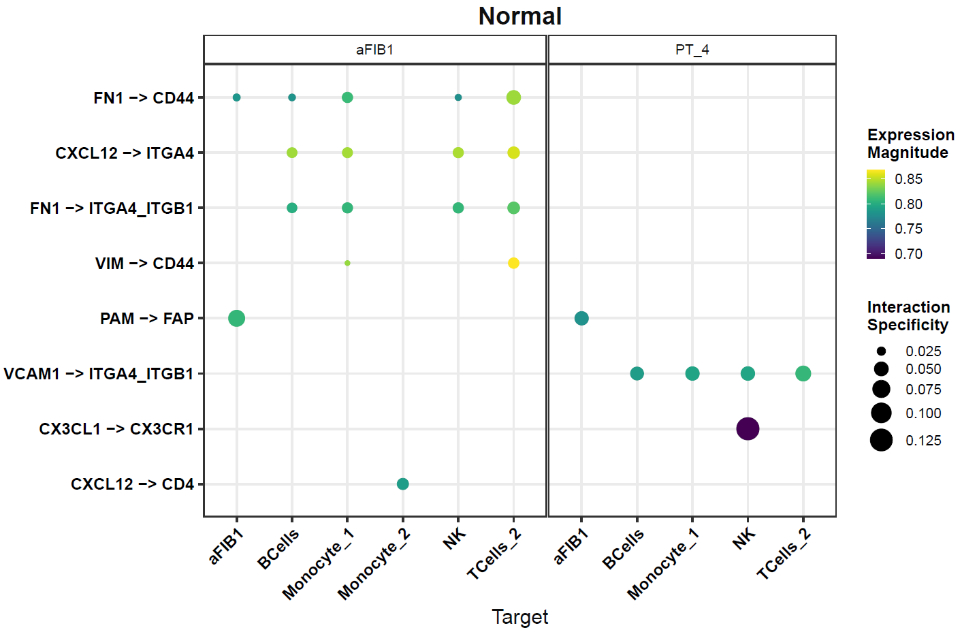

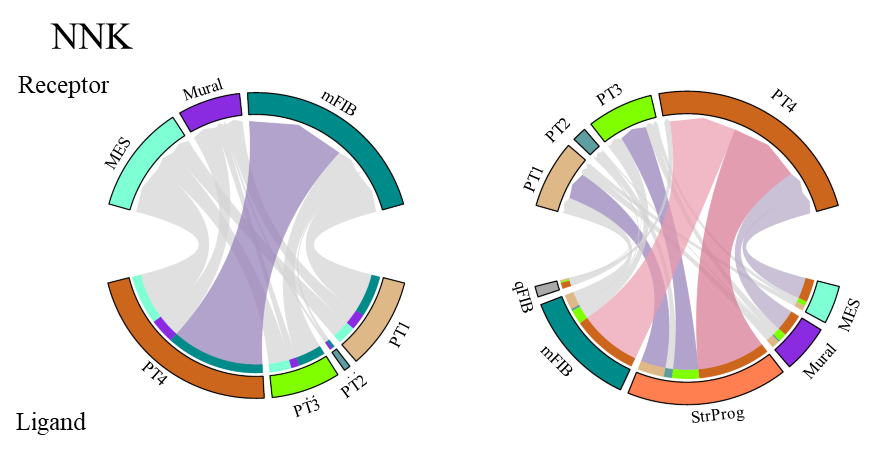

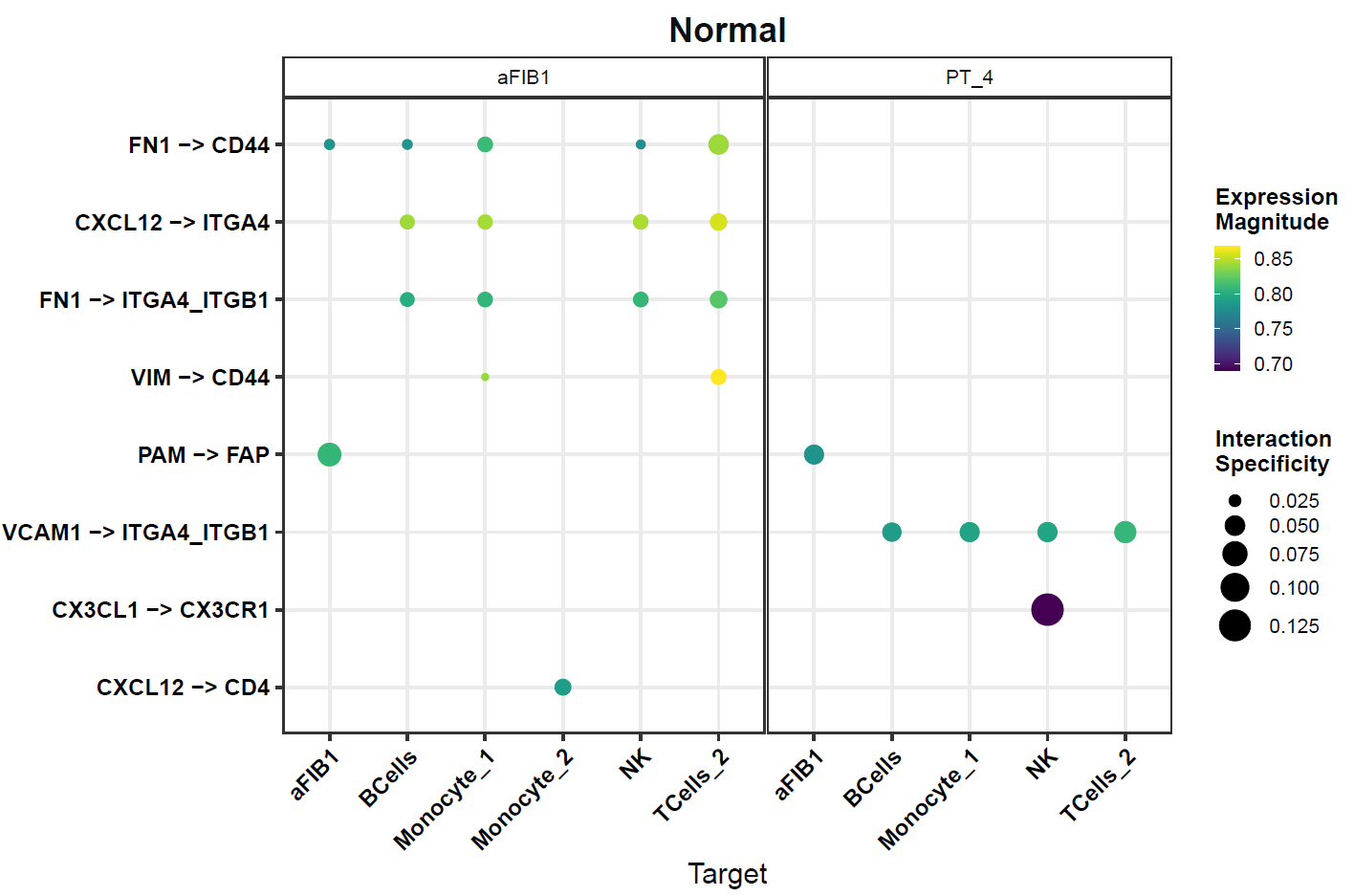

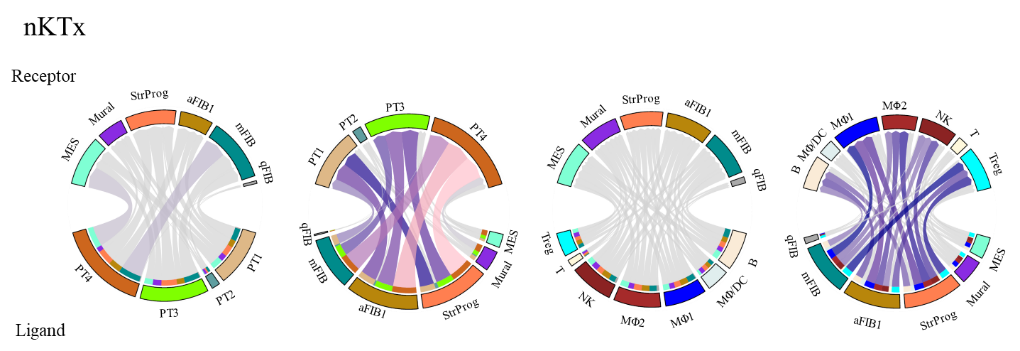
